# Haematopoietic loss of KDM6A impairs cardiac recovery in heart failure *via* epigenetic reprogramming of myeloid cells

**DOI:** 10.64898/2025.12.01.691512

**Authors:** Emmanouil G. Solomonidis, Dennis Hecker, Mariana Shumliakivska, Julian Leberzammer, Ariane Fischer, Minh-Thuy Katschke, Melanie Bendel, Lara Korth, Bianca Schuhmacher, Laura Rumpf, Simone-Franziska Glaser, Katja Schmitz, Guillermo Luxán, Silke Frenz, Penelope Pennoyer, Franziska Ganß, Evelyn Ullrich, David John, Stefan Guenther, Katalin Pálfi, Lisa M. Weiss, Matthias S. Leisegang, Ralf P. Brandes, David M. Leistner, Andreas M. Zeiher, Wesley T. Abplanalp, Mario Looso, Marcel H. Schulz, Stefanie Dimmeler, Sebastian Cremer

## Abstract

Clonal haematopoiesis (CH) is recognized as a potent independent risk factor for cardiovascular disease (CVD). While mutations in common CH-associated genes, such as *DNMT3A* and *TET2*, have been extensively studied, the pathological roles of other CH mutations remain poorly understood. Among these is KDM6A (UTX), an X-linked histone demethylase recently found to be commonly mutated in patients with heart failure. The mechanistic implications of KDM6A mutations in cardiac dysfunction remain largely unknown. Here, using multi-omics profiling and functional characterisation of murine models and patient-derived data, we demonstrate that haematopoietic loss of KDM6A substantially impairs cardiac recovery following myocardial infarction (MI). KDM6A deficiency enhances systemic and cardiac inflammation, characterized by augmented myeloid cell infiltration into the infarcted murine heart. Single-cell chromatin accessibility and single-cell RNA sequencing analyses revealed profound epigenetic and transcriptional reprogramming in KDM6A-deficient myeloid cells, notably CCR2⁺ recruited macrophages and neutrophils. These cells exhibited heightened inflammatory (*Il1b*, *Nlpr3*, *Saa3*) and chemotactic signatures (*Ccr2*, *Mif*, *Cxcl12*), increased activation of inflammatory transcription factor networks (AP-1, C/EBP), disrupted chromatin architecture, and enhanced glycolytic activity. Clinically, patients with heart failure harbouring KDM6A-driven CH exhibited increased pro-inflammatory monocyte signatures (*CCR2*, *NLPR3*, *NFKB1*, *FOS*, *JUN*, *IL6R*, *IL32*), underscoring the translational relevance. Integrative analyses further predicted pathogenic crosstalk between KDM6A-mutated monocytes and cardiac resident cells and was experimentally validated by demonstrating that KDM6A-silenced macrophages drive cardiomyocyte hypertrophy and cardiac fibroblast activation. Our findings establish a critical mechanistic link between KDM6A-driven CH, immune dysregulation, and worsened cardiac outcomes post-MI, highlighting novel avenues for personalized therapeutic strategies in heart failure.

## Introduction

Clonal haematopoiesis (CH) is increasingly recognized as an independent risk factor for cardiovascular disease (CVD)^1^. While mutations in commonly studied genes such as *DNMT3A* and *TET2* have been robustly linked to worse cardiovascular outcomes, many less frequently mutated genes remain poorly characterized. Recent large-scale analyses continue to identify novel and understudied CH-associated mutations that may promote disease through distinct mechanisms, emphasizing the potential for personalized medicine strategies tailored to specific genetic profiles^2^.

Among these understudied mutated genes is lysine demethylase 6A (KDM6A, also known as UTX), an X-linked histone demethylase^3^. The primary role of KDM6A is demethylation of H3 lysine 27 di- and tri-methylation (H3K27me2/H3K27me3), marks commonly associated with transcriptional repression^4^. UTY is the Y chromosome homolog of KDM6A, but lacks demethylase activity^5^. Recently, *KDM6A* emerged as one of the most commonly mutated genes among patients with heart failure, with mutations uniformly resulting in loss-of-function^6^. However, the mechanistic contribution of *KDM6A* mutations to cardiovascular pathology remains unexplored.

Here, we employed a comprehensive multi-omics approach in mouse models and patient-derived data to determine how haematopoietic loss of KDM6A function influences inflammation and cardiac dysfunction following ischemic injury. We reveal that haematopoietic KDM6A loss profoundly remodels chromatin and transcriptional programs in immune cells, resulting in augmented inflammatory responses and worsening cardiac function post-myocardial infarction (MI). These findings offer critical insights into CH-driven cardiovascular pathology and position heart failure patients harbouring KDM6A mutations as promising candidates for personalized therapeutic interventions targeting inflammation.

## Results

### Haematopoietic loss of KDM6A impairs recovery of cardiac function post-MI

To investigate how haematopoietic loss of KDM6A affects heart failure, we crossed *VaviCre* mice with *Kdm6a^fl/fl^* mice, generating mice with haematopoietic-specific Kdm6a knockout (*Kdm6a^Δ-Haem^*, **Figure 1A**). Baseline ejection fraction (EF) showed no differences between groups (**Figure 1B**). After MI induced by permanent occlusion of the left anterior descending (LAD) coronary artery, *Kdm6a^Δ-Haem^* mice displayed reduced EF compared to *Kdm6a^WT^* mice at day 28 post-MI (**Figure 1B**). Both male and female *Kdm6a^Δ-Haem^* mice exhibited impaired EF compared to *Kdm6a^WT^* controls at day 28 post-MI; however, the phenotype was more pronounced in females (**Supplementary Figure 1A**). Additionally, *Kdm6a^Δ-Haem^* hearts exhibited increased left ventricular scar size (**Figure 1C**) and increased expression of pro-inflammatory markers *Il1b* and *Nlrp3* at day 28 post-MI (**Figure 1D**). Analysis of the circulating leukocyte composition in peripheral blood revealed monocytosis at day 3 post-MI, the peak of inflammatory activity post-MI, specifically in female *Kdm6a^Δ-Haem^* mice, compared to controls (**Figure 1E-F**, **Supplementary Figure 1B**). T lymphocytes were reduced in both sexes at day 3 post-MI, while B cells were significantly decreased in females, with a similar trend observed in males (**Figure 1E-F**, **Supplementary Figure 1B**). Cardiac immune-cell profiling at day 3 post-MI revealed increased infiltration of myeloid cells, predominantly monocytes, into the ischemic myocardium (**Figure 1G-H**). At day 28 post-MI, histological staining confirmed an increased number of CD68⁺ macrophages in cardiac tissue (**Figure 1I-J**). Together, these data indicate that haematopoietic KDM6A loss enhances systemic inflammation, cardiac inflammation and myeloid cell recruitment into the myocardium, resulting in increased cardiac dysfunction after MI.

**Figure 1.**
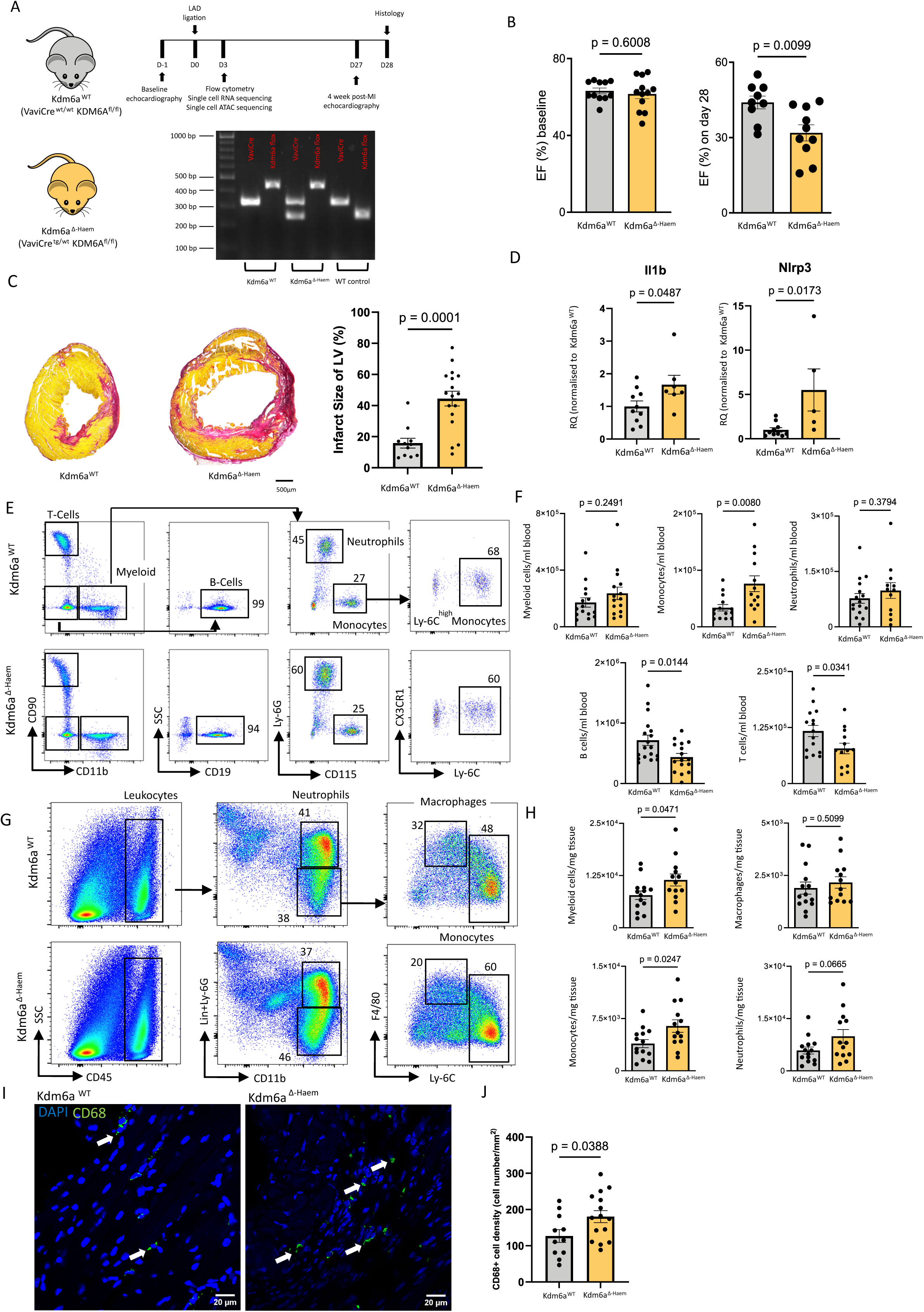
Haematopoietic loss of KDM6A impacts cardiac recovery while displaying pro-inflammatory features. (**A**) Experimental design of mouse studies with haematopoietic-specific KDM6A knockout. Agarose gel electrophoresis showing representative genotypes of *Kdm6a^WT^* (VaviCre^wt/wt^ KDM6A^fl/fl^), *Kdm6a^Δ-Haem^* (VaviCre^tg/wt^ KDM6A^fl/fl^) and wild type (WT) mice (VaviCre genotype: Internal positive control band 324 base pairs (bp), Cre transgene band 236 bp; KDM6A flox genotype: homozygous flox band 430 bp, WT band 249 bp) (**B**) Baseline cardiac ejection fraction (EF) (*Kdm6a^WT^* n=11 *vs Kdm6a^Δ-Haem^* n=12, p=0.601) and 28 days post-myocardial infarction (MI) (*Kdm6a^WT^* n=9 *vs Kdm6a^Δ- Haem^* n=10, p=0.001) (**C**) Representative images of Picro-Sirius Red cardiac tissue staining 28 days post-MI in *Kdm6a^WT^* and *Kdm6a^Δ-Haem^* mice, scale 500 μm. Quantification of scar size (*Kdm6a^WT^* n=11 *vs Kdm6a^Δ-Haem^* n=17, p=0.0001) (**D**) qPCR data from RNA extracted from cardiac tissue 28 days post-MI for *Il1b* (*Kdm6a^WT^* n=10 *vs Kdm6a^Δ-Haem^* n=7, p=0.04) and *Nlrp3* (*Kdm6a^WT^* n=10 *vs Kdm6a^Δ-Haem^* n=5, p=0.012) (**E**) Flow cytometry gating of blood leukocytes 3 days post-MI, numbers refer to % of parent gate (**F**) Flow cytometric analysis of blood leukocytes 3 days post-MI; CD11b+ myeloid cells (*Kdm6a^WT^* n=15 *vs Kdm6a^Δ-Haem^* n=15, p=0.249), monocytes (*Kdm6a^WT^* n=14 *vs Kdm6a^Δ-Haem^* n=13, p=0.008), B-cells (*Kdm6a^WT^* n=18 *vs Kdm6a^Δ-Haem^* n=16, p=0.014) and T-cells (*Kdm6a^WT^* n=15 *vs Kdm6a^Δ-Haem^* n=13, p=0.034) (**G**) Flow cytometry gating of cardiac leukocytes 3 days post-MI in *Kdm6a^WT^* and *Kdm6a^Δ-Haem^* mice, numbers refer to % of parent gate (**H**) Flow cytometric analysis of cardiac leukocytes 3 days post-MI; CD11b+ myeloid cells (*Kdm6a^WT^* n=14 *vs Kdm6a^Δ-Haem^* n=13, p=0.047), macrophages (*Kdm6a^WT^* n=14 *vs Kdm6a^Δ-Haem^* n=13, p=0.501), monocytes (*Kdm6a^WT^* n=14 *vs Kdm6a^Δ-Haem^* n=13, p=0.025) and neutrophils (*Kdm6a^WT^* n=14 *vs Kdm6a^Δ-Haem^* n=13, p=0.066). (**I**) Representative images of immunostaining of F4/80+ macrophages in cardiac tissue 28 days post-MI in *Kdm6a^WT^* and *Kdm6a^Δ-Haem^* mice. White arrows point to CD68+ macrophages. Scale bar: 20μm (**J**) Quantification of CD68+ macrophages 28 days post-MI in *Kdm6a^WT^* and *Kdm6a^Δ-Haem^* mice (*Kdm6a^WT^* n=11 *vs Kdm6a^Δ-Haem^* n=15, p= 0.039). Data are represented as mean ± SEM, p-values were calculated using two-tailed unpaired t-tests.

### KDM6A loss enhances inflammatory transcriptional and epigenetic activation in recruited macrophages post-MI

As KDM6A acts primarily as a transcriptional activator *via* H3K27me2/3 demethylation^3^, we next investigated how haematopoietic KDM6A loss affects the chromatin accessibility and expression signature of haematopoietic cells post-MI by performing single-cell assay for transposase-accessible chromatin sequencing (scATAC-seq) and single-cell RNA sequencing (scRNA-seq) on CD45⁺ immune cells isolated from cardiac tissue of *Kdm6a^Δ-Haem^* mice at 3 days post-MI. We selected female mice due to their more pronounced impairment in EF post-MI, increased systemic inflammation, and to avoid potential compensatory effects from Y chromosome homologues.

Following quality control (**Supplementary Figure 2A-B**) annotation of immune cell clusters was performed. Due to sparse data from scATAC-seq, we transferred cell-type labels from non-paired scRNA-seq data, based on marker gene expression (**Supplementary Figure 2C**). We cross-checked the successful annotation by checking the gene activity for markers used for scRNA-seq annotation, as assessed by the chromatin accessibility associated with the respective gene (**Supplementary Figure 2D**).

Clustering identified distinct immune subsets, including CCR2⁺ recruited macrophages, Ly6c-high monocytes, Ly6c-low monocytes, resident CCR2⁻ macrophages, interferon (Ifn)-activated monocytes, dendritic cells, neutrophils, T cells, and B cells (**Figure 2A-B**). Consistent with our flow cytometry data, *Kdm6a^Δ-Haem^* mice displayed increased frequency of myeloid cells, particularly CCR2⁺ macrophages and their Ly6c-high monocyte precursors (**Figure 2C**).

**Figure 2.**
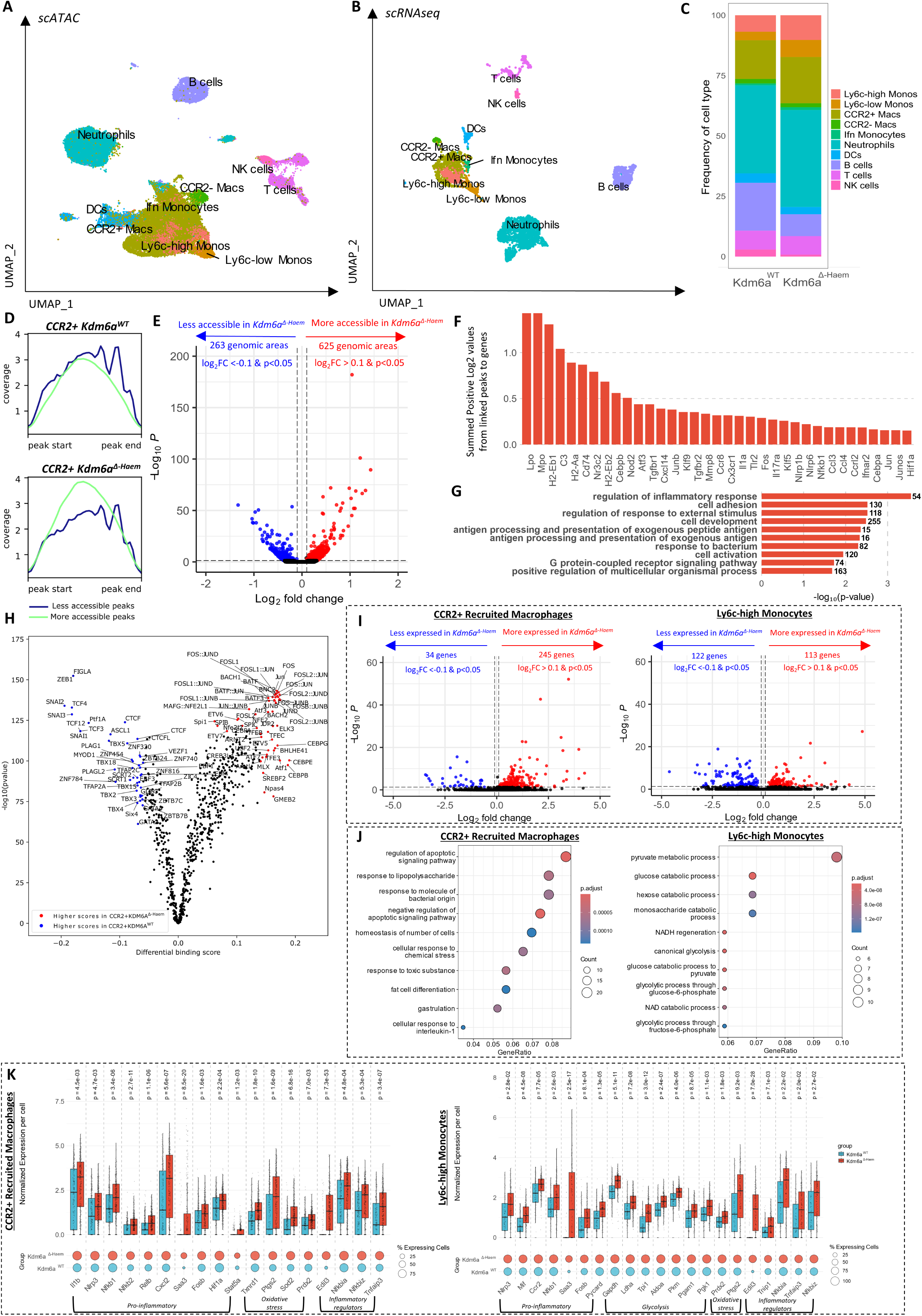
KDM6A loss enhances inflammatory transcriptional and epigenetic activation in recruited macrophages post-MI. (**A**) Uniform manifold approximation and projection (UMAP) clustering of single-cell ATAC sequencing data from cardiac CD45+ cells 3 days post-myocardial infarction in *Kdm6a^WT^* and *Kdm6a^Δ-Haem^* mice (n=3 female mice/group) (**B**) Uniform manifold approximation and projection (UMAP) clustering of single-cell RNA sequencing data from cardiac CD45+ cells 3 days post-myocardial infarction in *Kdm6a^WT^* and *Kdm6a^Δ-Haem^* mice (n=3 female mice/group) (**C**) Cell type distribution in *Kdm6a^WT^* and *Kdm6a^Δ-Haem^* cell clusters from single-cell RNA sequencing data (**D**) ATAC-seq coverage plots showing average chromatin accessibility signal (average counts per million) at genomic regions identified as differentially accessible peaks in CCR2⁺ macrophages from *Kdm6a^WT^* (top) and *Kdm6a^Δ-Haem^* (bottom) mice (**E**) Volcano plot of differentially accessible peaks in *Kdm6a^WT^* and *Kdm6a^Δ-Haem^* CCR2+ recruited macrophages (peak cut-off; 10% detection in either *Kdm6a^WT^* or *Kdm6a^Δ-Haem^* group) (**F**) Summed positive log2 fold-change values from differentially accessible chromatin peaks linked to genes in *Kdm6a^Δ-Haem^* CCR2+ recruited macrophages (**G**) Top enriched gene ontology (GO) terms enriched from enhancers linked to genes in *Kdm6a^Δ-Haem^* CCR2+ recruited macrophages. Numbers next to bars indicate the number of genes enriched in each GO (**H**) Differential transcription factor (TF) binding activity predicted with the tool TOBIAS from chromatin accessibility data in CCR2⁺ macrophages from *Kdm6a^Δ-Haem^* compared to *Kdm6a^WT^* mice. (**I**) Volcano plots of differentially expressed genes in CCR2+ recruited macrophages and Ly6c-high monocytes (gene cut-off; 10% expression in either *Kdm6a^WT^* or *Kdm6a^Δ-Haem^* group) (**J**) Top enriched GO terms enriched in differentially upregulated genes in *Kdm6a^Δ-Haem^* CCR2+ macrophages and *Kdm6a^Δ-Haem^* Ly6c-high monocytes (**K**) Normalized gene expression per cell (top) and the percentage of expressing cells (bottom) for differentially upregulated genes in CCR2⁺ recruited macrophages (left) and Ly6c-high monocytes (right) from *Kdm6a^Δ-Haem^* versus *Kdm6a^WT^* mice. Differentially expressed genes were identified using the Wilcoxon rank sum test, with Bonferroni correction for multiple comparisons.

Differential accessibility analysis revealed a significant number of differentially accessible genomic areas/peaks across all cell types with top affected clusters being CCR2+ macrophages and neutrophils. Interestingly, the majority of differential peaks were unique across cell types, suggesting that the loss of KDM6A has a unique impact in the different haematopoietic lineages (**Supplementary Figure 2F**).

Firstly, our analysis focused on CCR2⁺ recruited macrophages due to their increased infiltration into the infarcted myocardium. Aggregate ATAC-seq coverage plots revealed that in *Kdm6a^Δ-Haem^* CCR2⁺ macrophages, genomic regions diverged in accessibility compared to controls, reflecting a bidirectional shift in chromatin state (**Figure 2D**). More specifically, we identified more regions with increased accessibility (625) than with decreased accessibility (263) in *Kdm6a^Δ-Haem^* CCR2+ macrophages (**Figure 2E**), suggesting a tendency toward chromatin opening. This imbalance in accessibility suggests a shift to a potentially more transcriptionally active state in the absence of KDM6A.

To assess the potential functional relevance of the observed chromatin accessibility changes, we linked differentially accessible peaks to genes based on predicted peak-gene interactions. This analysis revealed that genes involved in inflammation and myeloid cell activation (*Il1a*, *Nfkb1*, *Cd74*, *Fos*, *Junb, Cebpa, Cebpb*) were most strongly associated with regions of increased accessibility in *Kdm6a^Δ-Haem^* CCR2⁺ macrophages (**Figure 2F**). Gene ontology (GO) enrichment analysis of significantly more accessible genes further supported this pro-inflammatory bias, with top enriched terms including “regulation of inflammatory response”, “cell adhesion,” and “response to external stimulus” (**Figure 2G**). These findings suggest that the epigenetic alterations observed in KDM6A deficient CCR2⁺ macrophages may prime macrophages for heightened inflammatory activation post-MI.

Chromatin accessibility changes additionally corresponded with altered transcription factor binding dynamics, notably increased binding of transcription factors from the AP-1 family, composed of various Jun (Jun, Junb, Jund), Fos (Fos, Fosb, Fosl1, Fosl2), and Atf subunits (Atf1, Atf3, Atf4, Batf), as well as members of the C/EBP family (e.g., Cebpa, Cebpb, Cebpe, Cebpg). Concurrently, we observed reduced binding of chromatin structural proteins such as CTCF, indicating potential disruption of chromatin architecture (**Figure 2H**).

We subsequently analyzed the downstream impact of the observed epigenetic changes using scRNAseq in CCR2⁺ recruited macrophages and Ly6c-high monocytes. In CCR2⁺ recruited macrophages, KDM6A deficiency led to a robust pro-inflammatory transcriptional response, with 245 genes upregulated and 34 downregulated compared to wild-type (**Figure 2I**) as GO term enrichment of upregulated genes revealed activation of inflammatory pathways (**Figure 2J**). *Kdm6a^Δ-Haem^* macrophages exhibited elevated expression of inflammatory mediators (*Il1b*, *Nlrp3*, *Cxcl2*, *Saa3*, *Fos*) alongside oxidative stress-related genes such as *Hif1a*, *Sod2*, and *Txnip* (**Figure 2K**), indicating a heightened inflammatory and oxidative state. Importantly, regulatory genes such as *Nfkbia*, *Tnfaip3*, and *Edil3* were also upregulated, suggesting the presence of compensatory mechanisms attempting to counterbalance the inflammatory activation.

In Ly6c-high monocytes, precursors of differentiated macrophages, 113 genes were upregulated and 122 downregulated in *Kdm6a^Δ-Haem^* cells (**Figure 2I**), indicating broader but, more moderate transcriptional reprogramming. GO analysis terms of enriched genes were dominated by glycolytic and carbohydrate catabolic processes (**Figure 2J**), which was reflected in the upregulation of glycolysis-related genes (*Gapdh*, *Pkm*, *Aldoa*, *Pgk1*, *Pgam1*). This metabolic shift occurred alongside increased expression of inflammatory genes (*Il1b*, *Ccr2*, *S100a8*, *Hif1a*) and oxidative stress markers (*Sod2*, *Txnip*), suggesting that KDM6A loss promotes an inflammatory phenotype that is both metabolically and oxidatively activated.

To better understand the upstream regulators driving these transcriptional changes, we applied LISA (Epigenetic Landscape In-Silico deletion Analysis)^7^. LISA predicted transcription factors and chromatin regulators responsible for the observed upregulated transcriptional signatures, highlighting significant enrichment for NF-κB signaling (RELA, CREBBP) and inflammatory activation (JUNB, CEBPA, CEBPB, STAT5A, STAT5B, JUN) in *Kdm6a^Δ-Haem^* recruited macrophages, consistent with previous transcription factor binding analysis (**Supplementary Figure 3**).

Collectively, these results demonstrate that KDM6A loss in recruited macrophages in MI induces extensive transcriptional and epigenetic reprogramming, marked by heightened inflammatory activation, disrupted chromatin architecture, and a metabolic shift toward increased glycolysis.

### KDM6A deletion induces transcriptional and chromatin reprogramming toward increased inflammatory and chemotactic signatures in neutrophils post-MI

Neutrophils represented another myeloid subset significantly altered transcriptionally and epigenetically in *Kdm6a^Δ-Haem^* mice following MI (**Supplementary Figure 2E-F**). Neutrophils exhibited similar bidirectional chromatin remodeling to CCR2^+^ macrophages upon KDM6A loss. However, in contrast to CCR2⁺ macrophages, most differentially accessible regions became less accessible (1030 down vs. 393 up; **Figure 3A-B**). Regions with increased accessibility were associated to genes related to enhanced activation of neutrophils (*Mpo*, *Lpo*, *Nr3c2*, *Thbs1*, *Cebpb*, *Nod2*, *Adam8*, *Junb*, *Fos*, *Il1r2*, *Stat6*) and chemotaxis (*Ccl6*, *Ccr1*, *Ccr3*, *Cxcr6*, *Cxcl5*, *Itgb2*, *Cd44*) (**Figure 3C**).

**Figure 3.**
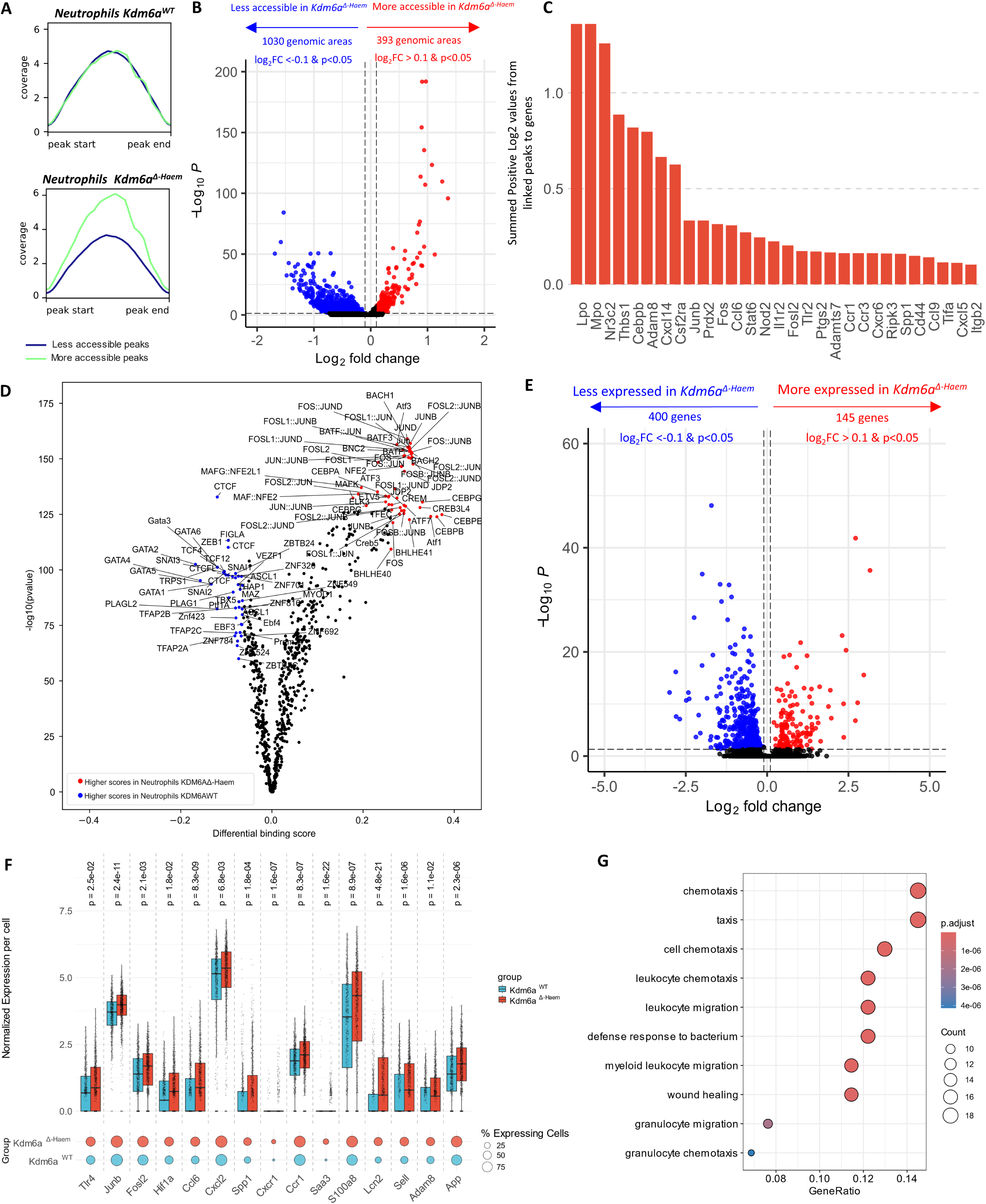
KDM6A deletion induces transcriptional and chromatin reprogramming toward increased inflammatory and chemotactic signatures in neutrophils post-MI. (**A**) ATAC-seq coverage plots showing average chromatin accessibility signal (average counts per million) at genomic regions identified as differentially accessible peaks in neutrophils from *Kdm6a^WT^* (top) and *Kdm6a^Δ-Haem^* (bottom) mice (**B**) Volcano plot of differentially accessible peaks in *Kdm6a^WT^* and *Kdm6a^Δ-Haem^* neutrophils (peak cut-off; 10% detection in either *Kdm6a^WT^* or *Kdm6a^Δ-Haem^* group) (**C**) Summed log2 positive fold-change values from differentially accessible chromatin peaks linked to genes in *Kdm6a^Δ- Haem^* neutrophils (**D**) Differential transcription factor (TF) binding activity predicted from chromatin accessibility data with the tool TOBIAS in neutrophils from *Kdm6a^Δ-Haem^* compared to *Kdm6a^WT^* mice (**E**) Volcano plots of differentially expressed genes in neutrophils (gene cut-off; 10% expression in either *Kdm6a^WT^* or *Kdm6a^Δ-Haem^* group (**F**) Normalized gene expression per cell (top) and the percentage of expressing cells (bottom) for differentially upregulated genes in neutrophils from *Kdm6a^Δ-Haem^* versus *Kdm6a^WT^* mice (**G**) Top enriched gene ontology (GO) terms enriched in differentially upregulated genes in *Kdm6a^Δ-Haem^* neutrophils. Differentially expressed genes were identified using the Wilcoxon rank sum test, with Bonferroni correction for multiple comparisons.

Corresponding shifts in chromatin accessibility aligned with altered transcription factor occupancy, including enhanced binding by AP-1 and C/EBP family members (**Figure 3D**), mirroring the trends observed in CCR2^+^ recruited macrophages. Interestingly, we observed diminished binding of GATA transcription factors (GATA1–5), a pattern previously linked to reduced anti-inflammatory regulation and promotion of pro-inflammatory myeloid phenotypes.^8,9^

Analysis of the genomic distribution of differentially accessible regions—covering promoters, exons, introns, intergenic, and untranslated regions (UTRs)—revealed widespread chromatin alterations across all cell types, indicating a broad disruption of chromatin structure, but with clear cell-type-specific patterns (**Supplementary Figure 4**). Notably, neutrophils exhibited prominent differential accessibility in promoters (34.64%), along with substantial changes in intronic (30.88%) and intergenic (26.18%) regions. In contrast, CCR2^+^ macrophages predominantly showed changes within intronic (44.99%) and intergenic (28.11%) regions (**Supplementary Figure 4**).

To assess the transcriptional consequences of chromatin remodeling in KDM6A-deficient neutrophils, we next performed differential gene expression analysis. A total of 145 genes were significantly upregulated, while 400 were downregulated in *Kdm6a^Δ-Haem^* neutrophils compared to control (**Figure 3E**). Many of the upregulated genes were associated with chemotaxis and immune activation, including *Saa3*, *Junb*, *Ccl6*, *Cxcr1*, *Ccr1*, *S100a8*, *Sell*, and *Cxcl2* (**Figure 3F**). GO term analysis of upregulated genes confirmed enrichment of pathways related to leukocyte chemotaxis and myeloid cell migration (**Figure 3G**).

Collectively, these data indicate that despite a global trend toward chromatin closure, KDM6A-deficient neutrophils selectively upregulate a focused inflammatory and migratory program, which likely contributes to their pathological role in sustained cardiac inflammation post-MI.

### Dysregulated monocyte transcriptomic signatures in heart failure patients with KDM6A-driven clonal haematopoiesis

To evaluate if our murine findings translate into clinical relevance, we profiled peripheral blood mononuclear cells (PBMCs) from patients with ischemic heart failure and reduced ejection fraction patients (HFrEF) with or without KDM6A-driven CH using scRNA-seq (**Figure 4A**). Patient characteristics are depicted in **Table 1**. UMAP clustering identified major immune subsets, including monocytes, T and B cells, mucosal-associated invariant T (MAIT) cells, plasmacytoid dendritic cells (pDCs), and myeloid dendritic cells (mDCs) (**Figure 4B-C**, **Supplementary Figure 5A-B**).

**Figure 4.**
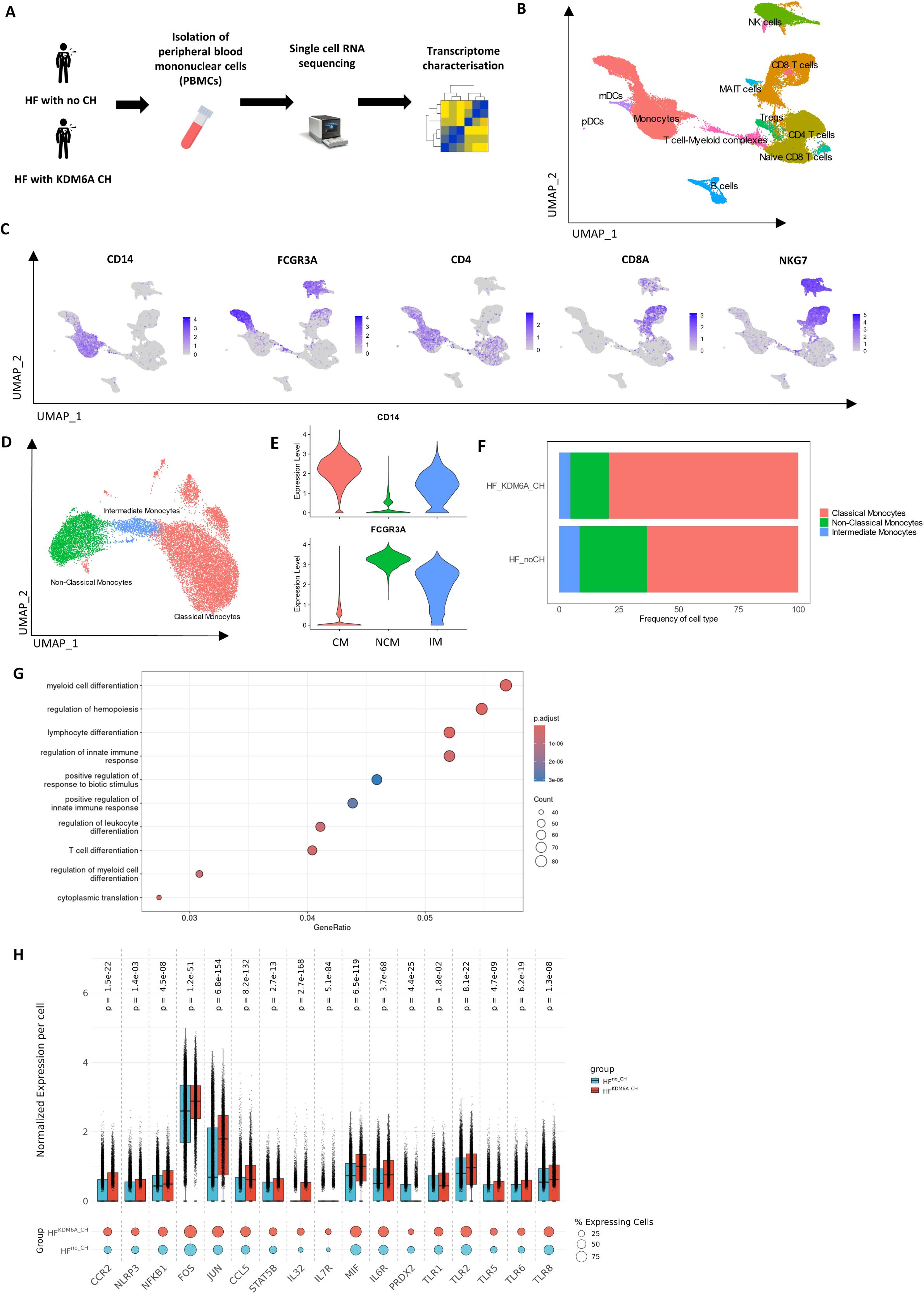
Dysregulated transcriptomic phenotype in monocytes from KDM6A CH heart failure. (**A**) Experimental design (**B**) Uniform manifold approximation and projection (UMAP) clustering of single-cell RNA sequencing data of PBMCs from heart failure (HF) patients with/without KDM6A clonal haematopoiesis (CH) (n=3 HF patients with no CH, n=2 HF patients with KDM6A CH) (**C**) UMAP plots of key peripheral leukocyte markers (**D**) UMAP of sub-clustered monocytes (**E**) Gene expression of *CD14* and *FCG3RA* in monocyte sub-clusters (**F**) Distribution of monocyte subtypes in HF patients with and without KDM6A CH (**G**) Top enriched gene ontology (GO) terms enriched in differentially upregulated genes in monocytes of HF patients with KDM6A CH (**H**) Normalized gene expression per cell (top) and the percentage of expressing cells (bottom) for differentially upregulated genes in neutrophils from HF*^KDM6A_CH^* versus HF*^no_CH^* patients. Differentially expressed genes were identified using the Wilcoxon rank sum test, with Bonferroni correction for multiple comparisons. Differential gene fold cuttoffs for GO term analysis was log_2_FC≥±0.1, adjusted p- value < 0.05 and at least 10% gene expression in either group.

**Table 1.** Patient characteristics. Patient characteristics of heart failure patients with and without KDM6A driven clonal haematopoiesis.

| Characteristic | HF_Control_1 | HF_Control_2 | HF_Control_3 | HF_KDM6A_1 | HF_KDM6A_2 |
| --- | --- | --- | --- | --- | --- |
| Sex | male | male | male | male | male |
| nt pro BNP | 756.7 | 1253 | 1235 | 6402 | 2779 |
| EF | 25 | 25-30 | 40 | 30 | 40 |
| Age | 61 | 79 | 68 | 84 | 81 |
| Hypertension | N | N | Y | Y | N |
| Kidney disease | Y | Y | N | Y | N |
| Hyperchol | Y | Y | Y | Y | Y |
| Statin | Y | Y | Y | Y | Y |
| ACE/ARB/ARNI | Y | Y | Y | Y | Y |
| BB | Y | Y | Y | Y | Y |
| Vessel | 3 | 3 | 3 | 3 | 2 |
| Leuk/nl | 6.75 | 5.87 | 9.87 | 6.6 | 8.23 |
| Trop | 28 | 29 | 13 | 26 | 30 |
| NYHA | 3 | 3 | 3 | 3 | 2 |
| Diabetes | N | Y | Y | Y | N |
| CRP | N/A | 0.72 | 0.17 | 0.52 | 0.18 |
| VAF KDM6A (%) | N/A | N/A | N/A | 17.2 | 2.59 |
| KDM6A Mutation info DNA | N/A | N/A | N/A | 824_825delAGinsT | c.3133G>A |
| KDM6A Mutation info Protein | N/A | N/A | N/A | K275fs | p.Glu1045Lys |

Differential gene expression analysis revealed transcriptional differences across multiple leukocyte subsets (**Supplementary Figure 5C**). Given the myeloid-specific inflammatory phenotype observed in *Kdm6a^Δ-Haem^* mice, we focused our analysis on monocyte populations. Sub-clustering revealed classical (CD14⁺), intermediate, and non-classical (CD16⁺) monocytes (**Figure 4D-E**). Notably, HF patients with KDM6A-CH showed a marked expansion of classical monocytes, the subset most associated with inflammatory responses (**Figure 4F**). Differential expression analysis across all monocyte subsets identified upregulated genes in KDM6A-driven CH involved in myeloid differentiation, immune regulation, and leukocyte activation (**Figure 4G**). This included genes such as *CCR2*, *NFKB1*, *FOS*, *JUN*, *CCL5*, *STAT5B*, *IL32*, *IL7R*, *MIF*, and multiple Toll-like receptors (*TLR1*, *TLR2*, *TLR5*, *TLR6*, *TLR8*), closely mirroring the inflammatory signatures observed in mouse *Kdm6a^Δ-Haem^* myeloid cells (**Figure 4H**). These results suggest that KDM6A-CH in human HF patients is associated with enhanced myeloid-driven inflammation and may contribute to disease progression.

### Altered interactions in the cardiac milieu driven by KDM6A CH monocytes

To understand how KDM6A-mutated monocytes influence cardiac tissue in heart failure, we performed CellChat^10^ analysis by integrating monocyte scRNA-seq data from HF patients with and without KDM6A-CH and publicly available heart single-nucleus RNA-seq (snRNA-seq) from patients with HFrEF^11^ (**Figure 5A-D, Supplementary Figure 6A-C**) (patient details published previously^12^). Cell-cell interaction modelling revealed a significant increase in inferred outgoing signalling from KDM6A-CH monocytes toward multiple cardiac cell populations, including fibroblasts and cardiomyocytes (**Figure 5E**). Among the top enriched pathways towards fibroblasts and cardiomyocytes were IL-6, CCL, HGF, and MHC-I/II signalling, implicating both inflammatory and fibrotic communication circuits (**Figure 5F**). To functionally validate these predictions, we silenced KDM6A in human monocyte-derived macrophages using siRNA and exposed primary cardiomyocytes and cardiac fibroblasts to their conditioned media (**Figure 5G-H**). Cardiomyocytes treated with KDM6A-deficient macrophage supernatant exhibited significantly increased beating frequency and hypertrophy (**Figure 5I-J**). Similarly, fibroblasts exposed to the same conditioned medium showed elevated collagen alpha-1(I) chain production, indicating enhanced fibrotic activation (**Figure 5K**). These findings suggest that monocytes harbouring KDM6A mutations may exacerbate HF pathology by aberrantly reshaping paracrine signalling in the cardiac microenvironment.

**Figure 5.**
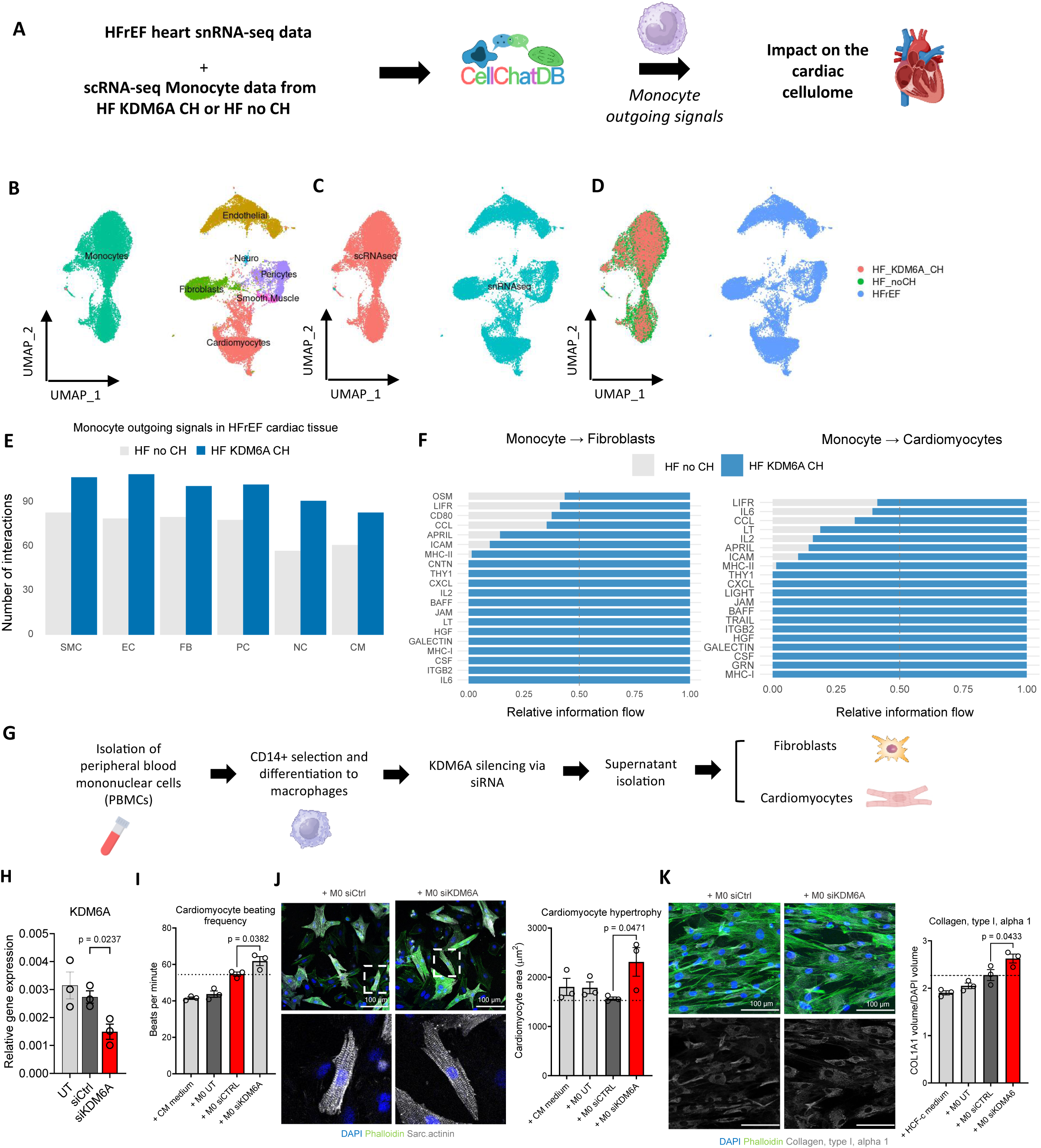
Altered interactions in the cardiac milieu driven by KDM6A CH monocytes (A) Analysis workflow. (**B**) UMAP plots of integrated single-cell RNA-seq of HF patients with/without KDM6A clonal haematopoiesis (CH) and single-nucleus RNA-seq of cardiac cells from n=3 HF patients with reduced ejection fraction (HFrEF), here grouped by cell type, (**C**) grouped by sequencing type, (**D**) grouped by group (**E**) Outgoing signals from monocyte with/without KDM6A CH towards HFrEF cardiac cells, as inferred by CellChat (FB; fibroblasts, EC; endothelial cells, SMC; smooth muscle cells, PC; pericytes, NC; neuronal cells, CM; cardiomyocytes) (**F**) Enriched signalling pathways in monocytes with KDM6A CH towards HFrEF cardiac fibroblasts and cardiomyocytes (**G**) in-vitro experimental design (**H**) siRNA knockdown of KDM6A in human monocyte-derived M0 macrophages (relative gene expression, normalised to *RPLP0*, p=0.024) (**I**) Rat cardiomyocyte beating frequency upon stimulation with supernatant from KDM6A-silenced M0 macrophages (p=0.038) (**J**) Representative images of rat cardiomyocytes stained with sarcomeric actinin, phalloidin and DAPI. Zoomed regions are derived from white boxed areas. Scale bar: 100μm. Quantification of cardiomyocyte area (p=0.047) (**K**) Human cardiac fibroblast activation assay as measured by Collagen type I alpha 1 immunostaining in fibroblasts treated with supernatant from KDM6A-silenced M0 macrophages (p=0.043), and representative images, scale bar: 100μm. Data are represented as mean ± SEM, p-values were calculated using two-tailed unpaired t-tests or one-way ANOVA.

## Discussion

In this study we demonstrate for the first time that lack of haematopoietic KDM6A is detrimental in the context of HF. We show that loss of KDM6A function in haematopoietic cells results in worsened cardiac function after MI through an enhanced inflammatory activation of myeloid cells as shown by increased production of pro-inflammatory cytokines. Haematopoietic KDM6A deletion results in a chromatin landscape rewiring that promotes binding of TFs such as AP-1 that drive expression of pro-inflammatory programs. These findings are corroborated by human heart failure patient data, where heart failure patients with mutated KDM6A-driven CH show enhanced inflammatory signalling in circulatory monocytes with a potential to aberrantly activate cardiac cells such as fibroblasts and cardiomyocytes upon infiltration, reinforcing the link between KDM6A mutations and worsened cardiovascular outcomes.

Additionally, haematopoietic deficiency of KDM6A results in worse EF following MI, which is associated with increased recruitment of myeloid cells to the cardiac tissue. These recruited myeloid cells exhibit a pronounced pro-inflammatory phenotype, as evidenced by the upregulation of canonical inflammatory and pro-recruitment markers, including IL1B, NLRP3, and CCR2, all of which are associated with adverse cardiovascular outcomes^13–15^. Furthermore, KDM6A deficient immune cells display dysregulated communication towards other cardiac cell types, which may further drive attenuated cardiac healing post-MI. In-depth analysis of the inflammatory profile of KDM6A-deficient myeloid cells reveals concurrent upregulation of transcription factors from the NF-κB and AP-1 families, including JUN and FOS, which act upstream of classical pro-inflammatory cytokines^16^ and are linked to aggravated cardiac inflammatory response and poor cardiac remodelling post-MI^17^. In parallel, these cells display a metabolic shift towards enhanced glycolysis, a hallmark feature of myeloid cell inflammatory activation^18^. Interestingly, despite the heightened pro-inflammatory response, KDM6A-deficient myeloid cells also exhibit increased expression of anti-inflammatory genes, including NF-κB regulatory elements and EDIL3 (also known as DEL-1), a molecule implicated in the resolution of inflammation^19^. This suggests an intrinsic, yet dysregulated, attempt by these cells to counterbalance their exacerbated immune activation.

Moreover, chromatin accessibility profiling of KDM6A-deficient myeloid cells reveals profound alterations in chromatin structure. A substantial number of genomic regions exhibit increased accessibility, while others were repressed, collectively resulting in an overall chromatin landscape rewiring that favours the binding of pro-inflammatory transcription factors such as members of the AP-1 and C/EPB family. Given that KDM6A is classically recognized as a transcriptional activator, these findings appear counterintuitive. However, they underscore the complex regulatory role of KDM6A in immune cell chromatin remodelling following MI, suggesting potential secondary effects mediated by other chromatin-modifying factors in its absence. Notably, KDM6A-deficient CCR2⁺ recruited macrophages exhibit a greater proportion of chromatin opening compared to neutrophils, which show more repressed genomics areas. Additionally, although the majority of differentially accessible regions are unique to each cell type, indicating that KDM6A loss remodels the chromatin landscape in a lineage-specific manner, both KDM6A deficient CCR2⁺ macrophages and neutrophils act *via* similar transcription factor programs, particularly AP-1 and C/EBP. This convergence in upstream regulatory networks is accompanied by a shared biological outcome of heightened expression of inflammatory and chemotactic genes.

Histone demethylases are important in immunity^20^ and implicated in the regulation of cardiovascular disease^21^. KDM6A is associated with inflammation, albeit not in the context of cardiovascular disease. KDM6A deficiency in the urothelium led to activated cytokine and chemokine pathways in the context of TP53-induced bladder cancer^22^. However, KDM6A contributes to the innate response during inflammation by promoting IL-6 and IFN-β production in LPS-treated mouse peritoneal macrophages^23^. These contrasting observations suggest that the role of KDM6A during inflammation may differ depending on context, severity, or the specific cell type involved, and may further diverge when comparing partial loss versus complete ablation of KDM6A function. Recently, KDM6A was found to regulate immune responses in the context of multiple myeloma by binding and regulating genes associated with immune recognition and cytokine signalling such as NLRC5 and CIITA^24^. Moreover, KDM6A and KDM7A bind the NF-κB binding site in TNF-α-induced endothelial cells, which suggests that removal of repressive histone marks by KDM6A is critical for NF-κB-dependent regulation of genes that control inflammation^25^. When using an inhibitor against KDM6A (GSK-J1), it has been shown that there is a reduction in lipopolysaccharide-induced proinflammatory cytokine production by human primary macrophages.^26^ However, there is conflicting evidence regarding the regulatory role of KDM6A in inflammation as certain studies have shown that upon inhibition of KDM6A there is improvement in diabetes-induced renal abnormalities in an animal model of type 2 diabetes by reducing inflammation^27^. In summary, KDM6A plays a significant role in immune responses and the regulation of inflammation, but the effects of KDM6A on inflammation are complex and context-dependent.

Furthermore, our human data provide important clarity by directly linking KDM6A-driven CH to pathogenic inflammation in heart failure. Monocytes from patients harbouring KDM6A mutations exhibit upregulation of key inflammatory mediators, including *IL1B*, *NFKB1*, *FOS*, and *JUN*, closely mirroring the transcriptional profiles seen in *Kdm6a^Δ-Haem^* mice. These shared signatures point to a conserved mechanism of immune activation and support a role for KDM6A in restraining myeloid-driven inflammation in the context of HF. Our findings suggest potential implications for personalized medicine, particularly regarding anti-cytokine therapies for patients with KDM6A-driven CH. This is supported by recent data from a follow-up analysis of the CANTOS^28^ randomized clinical trial, which suggested that MI patients harbouring TET2-driven CH could respond better to canakinumab, targeting IL-1β, compared to those without CH^29^.

Certain limitations should be considered when interpreting this study. Firstly, the relatively small patient cohort for scRNA-seq analysis may restrict the generalizability of our findings; larger, diverse cohorts would strengthen our conclusions. Moreover, while the *Kdm6a^Δ-Haem^* mouse model provided valuable mechanistic insight into KDM6A-driven inflammation in heart failure, it is important to acknowledge the clinical heterogeneity of KDM6A mutations. However, all previously reported mutations to date appear to result in loss-of-function, supporting the relevance of our knockout model^30–38^. Additionally, it remains unresolved why KDM6A loss promotes pro-inflammatory programs post-MI. These responses could involve genome-wide chromatin dysregulation affecting enhancer-promoter interactions, possibly mediated by alterations in histone modifications (H3K27me3, H3K4me1, H3K27ac). Future research addressing epigenetic priming in haematopoietic progenitors and targeted therapeutic interventions could clarify these mechanisms, further highlighting personalized strategies for patients with KDM6A-driven clonal haematopoiesis.

In summary, our findings reveal a pivotal role for KDM6A in orchestrating immune cell responses during heart failure after ΜΙ, highlighting the therapeutic potential of modulating inflammatory pathways to improve outcomes in patients with CH driven by KDM6A mutations.

## Materials & Methods

### Animals

All animal experiments were conducted according to the principles of laboratory animal care and according to German national laws. The studies were approved by the local ethics committee (Regierungspräsidium Darmstadt, Hessen) in application FU/1239. *VaviCre* (*B6.Cg-Commd10^Tg(Vav1-icre)A2Kio^/J,* Jackson laboratory) heterozygous male mice were bred with female *Kdm6a^fl/fl^* (*B6;129S-Kdm6a^tm1.1Kaig^/J,* Jackson laboratory) homozygous mice to establish *VaviCreKdm6a^fl/f^* offspring with a haematopoietic specific KDM6A knockout (referred to in this manuscript as *Kdm6a^Δ-Haem^* mice). Control mice (referred to in this manuscript as *Kdm6a^WT^* mice) were littermates lacking the *VaviCre* transgene. Phenotyping experiments included both male and female mice. Single cell RNA/ATAC experiments included female mice only.

### Genotyping

Genotyping was conducted with DNA isolated from ear punches using touchdown PCR according to the protocols provided by the Jackson laboratory for the respective mouse strains. Briefly, the presence of the *VaviCre* transgene was validated using the following primers: Internal Positive Control Forward; 5’-CTAGGCCACAGAATTGAAAGATCT-3’, Internal Positive Control Reverse; 5’-GTAGGTGGAAATTCTAGCATCATCC-3’, Transgene Forward; 5’-AGATGCCAGGACATCAGGAACCTG-3’, Transgene Reverse; 5’-ATCAGCCACACCAGACACAGAGATC-3’. The presence of *Kdm6a^fl/fl^* transgene was validated using the following primers: Forward; 5’-GCTACTGGGGTGTTTTGAATG-3’, Reverse; 5’-TTTCATAGAACAGTTTCAGGATACC-3’.

### Myocardial infarction model (left anterior descending coronary artery ligation)

The animals were sedated using isoflurane, and pain relief was provided through an intraperitoneal injection of Buprenorphine (0.1 mg/kg body weight) along with an intercostal nerve block using Bupivacaine (1 mg/kg body weight, 0.25% concentration). For postoperative pain management, Buprenorphine and Carprofen (5 mg/kg body weight) were administered every 12 or 24 hours for three days. To prevent infection after surgery, Ampicillin (100 mg/kg body weight) was added to the drinking water. The myocardial infarction (MI) was induced by permanently ligating the left anterior descending coronary artery while the animals were under mechanical ventilation.

### Echocardiography

Echocardiographic assessments were performed using the Vevo 3100 system (Fujifilm) under isoflurane anesthesia at baseline and day 28 post-MI. The acquired images were analyzed in a blinded manner with VevoLab 5.5.0 software. Both systolic and diastolic phases were evaluated by capturing images from the short and long axes of the ventricles to determine ejection fraction and left ventricular mass. The left atrial size was measured in the four-chamber view during the end-systolic phase with the mitral valve in the closed position. Diastolic function was assessed by examining the velocity of blood flow through the mitral valve using pulsed Doppler, while the mitral valve annulus motion was evaluated with tissue Doppler imaging. From these measurements, the E/A and E/e’ ratios were calculated.

### Murine tissue processing

Peripheral blood: Mouse blood was isolated from sacrificed mice from the vena cava following culling *via* cervical dislocation under isoflurane anaesthesia or sampled from live mice via the tail vein prior to MI. Red blood cells with lysed with 1× red blood cell lysis buffer (BioLegend) after which point samples were processed for flow cytometric analysis (See “Flow cytometry analysis of murine tissue” section).

Heart: Mice were sacrificed using cervical dislocation under isofluorane anaesthesia. Hearts were perfused *via* the left ventricle with cold PBS and excised before being dissected and enzymatically digested (in RPMI 1640 media; 450 U/ ml collagenase I, 125 U/ml collagenase XI, 60 U/ml DNase I, and 60 U/ml hyaluronidase, 30 minutes, 37°C degrees). Following digestion, mechanical dissociation was carried out (gentleMACS™) and cells were filtered through a 40-μm nylon mesh to obtain a single cell suspension.

### Flow cytometry analysis of murine tissue

Flow cytometry measurements were obtained using an LSRFortessa^TM^ X-20 (BD) with the following setup: UV laser (355 nm) with 379/28, 525 /50 filters; Violet laser (405nm) with 450/50, 525/50, 610/20, 670/80, 710/50, 780/60 filters; Blue laser (488 nm) with 530/30, 710/50 filters; Yellow/green laser (561 nm) with 586/15, 610/20, 670/30, 710/50, 780/60 filters; Red (640 nm) laser with 670/30, 730/45, 780/60 filters. Data analysis was performed with Flowjo v10.

Peripheral blood: Following washing of peripheral leukocytes, cells were stained with antibodies against Cd45 (Biolegend, clone 30-F11), Cd11b (Biolegend, clone M1/70), Cd90 (Biolegend, clone 30-H12), Ly6G (Biolegend, clone 1A8), Cd115 (Biolegend, clone AFS98), Cx3cr1 (Biolegend, clone SA011F11), Cd19 (BD Bioscience, clone 1D3) and Ly6C (Biolegend, clone HK1.4). Immune cell subsets were identified as follows: Monocytes (Cd45^high^, Cd11b^high^, Ly6G^low^, Cd115^high^, Cx3cr1^high^, Ly6C^low/high^), Neutrophils (Cd45^high^, Cd11b^high^, Ly6G^high^), T-cells (Cd45^high^ Cd11b^low^, Cd90^high^) and b-cells (Cd45^high^, Cd11b^low^, Cd19^high^).

Heart: Single-cell suspensions of murine hearts were stained with antibodies against Cd45 (Biolegend, clone 30-F11), Ly6c (Biolegend, clone HK1.4), F4/80 (Biolegend, clone BM8), Cd11b (Biolegend, clone M1/70) MHC2 (Biolegend, clone M5/114.15.2) and linage exclusion of Ly6G (Biolegend, clone RB6-8C5), Cd90 (Biolegend, clone 30-H12), Cd103 (Biolegend, clone 2E7), Ter119 (Biolegend, clone Ter119), B220 (Biolegend, clone RA3-6B2) and Nk-1.1 (Biolegend, PK136). Cardiac immune cells were defined as: Monocytes (Cd45^high^, Ly6G^low^, Ly6C^high^, F4/80^low^), Macrophages (Cd45^high^, Ly6G^low^, Ly6C^low^, F4/80^high^), Neutrophils (Cd45^high^, Cd11b^high^, Ly6G^high^), T-cells (Cd45^high^ Cd11b^low^, Cd90^high^) and B-cells (Cd45^high^, Cd11b^low^, Cd19^high^).

### Murine tissue histology & immunofluorescent staining

Hearts were fixed in 4% paraformaldehyde and subsequently embedded in paraffin. The tissue was then sectioned into 4 μm thick slices using a microtome. After deparaffinization, the slides were stained with Picro-Sirius Red to assess fibrosis. The slides were mounted with Prolong™ Gold Antifade Mountant (ThermoFisher Scientific), and images were captured with a Nikon Eclipse Ti2 microscope. For immunofluorescent staining of macrophages in the post-MI heart, a primary rabbit anti-mouse CD68 antibody (Cell Signaling, clone E307V) was used, along with a secondary polyclonal fluorochrome-conjugated donkey anti-rabbit antibody (Thermo Fisher Scientific). The slides were mounted with Prolong™ Gold Antifade Mountant (ThermoFisher Scientific), and imaging was performed on a Zeiss LSM 780 Observer Z.1 microscope. Image analysis was carried out using ZEN software and ImageJ. CD68+ cell density was calculated as the number of CD68+ cells per region of interest, normalized to area (cells/mm²), where each ROI was 212.55 × 212.55 µm.

### RNA isolation, cDNA synthesis and quantitative RT-qPCR

Total RNA was isolated using the RNeasy Plus Mini Kit (Qiagen) following the manufacturer’s guidelines. RNA content and purity were assessed by spectrophotometry (NanoDrop Technologies). mRNA expression was quantified by RT-qPCR using total RNA, which was reversed transcribed by MuLV reverse transcriptase (Thermo Fisher Scientific) and random hexamer primers (Thermo Fisher Scientific). cDNA synthesis was performed according to the protocol: (Primer annealing) 65°C x 5 min, (Primer elongation) 25°C x 10 min, (Reverse transcription) 37°C x 50 min, (Enzyme denaturation) 70°C x 15 min. Expression levels of mRNA were detected by using Fast SYBR Green Applied Biosystems) and an Applied Biosystems Viia7 machine. Cycling conditions on the machine were as follows: (Hold) 95°C x 30 sec, (PCR) 95°C x 1 sec, 60°C x 20 sec repeated for 40 cycles, (Melt curve) 95°C x 15 sec, 60°C x 60 sec, 95°C x 15 sec. Relative gene expression was calculated with the QuantStudio Real-Time PCR software (version 1.3) using 2^-ΔCt^ (ΔCt= Ct target gene – Ct housekeeping gene). RPLP0 was used as housekeeping gene.

Mouse primer sequences: *Rplp0*-F: 5’-TCGACAATGGCAGCATCTAC-3’, *Rplp0* R: 5’-ATCCGTCTCCACAGACAAGG-3’, *Il1b*-F: 5’-TGGAAAAGCGGTTTGTCT-3’, *Il1b*-R: 5’-ATAAATAGGTAAGTGGTTGCC-3’, *Nlrp3*-F: 5’-AGAAGAGACCACGGCAGAAG-3’, *Nlrp3*-R: 5’-CCTTGGACCAGGTTCAGTGT-3’. Human primer sequences were: *RPLP0*-F: 5’-ATCCGTCTCCACAGACAAGG-3’, *RPLP0*-R: 5’-TCGACAATGGCAGCATCTAC-3’, *KDM6A*-F: 5’-AGCGCAAAGGAGCCGTGGAAAA-3’, *KDM6A*-R: 5’-GTCGTTCACCATTAGGACCTGC-3’.

### Cardiac CD45+ cell isolation for single-cell RNA sequencing

Heart tissue processing was performed according to the section “Murine tissue processing” with a few alterations. After obtaining the cardiac single cell suspension a tissue debris removal was used according to manufacturer’s instructions (Miltenyi). Subsequently, red blood cells were lysed with 1x RBC lysis buffer (Biolegend) for 5 minutes. Following washing, cells were stained with Cd45 (Biolegend, clone 30-F11) and 7-AAD (BD) and sorting of CD45^high^, 7-ADD^Neg^ cells were performed for downstream single-cell analyses. Sorting was performed with an Aria Fusion (BD) with the following setup: UV laser (355 nm) with 379/28, 730 /45 filters; Violet laser (405nm) with 450/40, 525/50, 610/20, 660/20, 710/50, 780/60 filters; Blue laser (488 nm) with 488/10, 530/30, 584/42, 616/23, 695/40, 780/60 filters; Red (640 nm) laser with 670/30, 730/45, 780/60 filters.

### Nuclei isolation from cardiac CD45+ cells for single-cell ATAC sequencing

Murine cardiac tissue processing for CD45+ cell sorting was processed as described in the segment “Cardiac CD45+ cell isolation for single-cell RNA sequencing”. Nuclei were extracted with a modified version of 10x Genomics’s “Nuclei Isolation for Single Cell ATAC Sequencing” protocol. Briefly, CD45+ cells were centrifuged (300g, 5min, 4°C) and resuspended in pre-chilled lysis solution (Tris-HCl pH 7.4 10mM, NaCl 10mM, MgCl_2_ 3mM, Tween-20 0.1%, Nonidet P40 Substitute 0.1%, Digitonin 0.01%, 1% BSA) for 5 minutes on ice. Nuclei were then washed with a 1ml wash solution (Tris-HCl pH 7.4 10mM, NaCl 10mM, MgCl_2_ 3mM, Tween-20 0.1%, 1% BSA) and centrifuged (500g, 5min, 4°C). Viable nuclei were stained with 7-ADD and 7-ADD^high^ nuclei were FACS sorted before proceeding to downstream single-cell ATAC library preparation. Sorting was performed with an Aria Fusion (BD) (set up described in section “Cardiac CD45+ cell isolation for single-cell RNA sequencing”).

### Single-cell RNA sequencing

Viable murine CD45+ cell suspensions or human PBMCs were processed according to the manufacturer’s instructions, with cells loaded onto a 10X Chromium Controller (10X Genomics). The libraries for single-cell RNA sequencing (scRNA-seq) were prepared using the Chromium Single-Cell 3′ v3 Reagent Kit (10X Genomics). Initially, individual cells were isolated into droplets within an emulsion, each containing gel beads coated with unique primers that carried 10X cell barcodes, unique molecular identifiers, and poly(dT) sequences. Reverse transcription then generated barcoded full-length cDNA, which was followed by emulsion disruption using a recovery agent and cDNA purification with DynaBeads MyOne Silane Beads (Thermo Fisher Scientific). Amplification of cDNA was performed on a Biometra Thermocycler TProfessional Basic Gradient with a 96-Well Sample Block under the following conditions: 98 °C for 3 minutes, followed by 14 cycles of 98°C for 15 seconds, 67°C for 20 seconds, and 72°C for 1 minute, ending with a 72°C step for 1 minute and holding at 4°C. The amplified cDNA product was purified using the SPRIselect Reagent Kit (Beckman Coulter). To construct the indexed sequencing libraries, the Chromium Single-Cell 3′ v3 Reagent Kit was used to carry out fragmentation, end repair, A-tailing, and size selection with SPRIselect. Further steps included adaptor ligation, post-ligation cleanup with SPRIselect, polymerase chain reaction for sample index, and cleanup using SPRIselect beads. Quantification and quality assessment of the libraries were done using the Bioanalyzer Agilent 2100 with a High-Sensitivity DNA chip (Agilent Genomics). The indexed libraries were pooled equimolarly and sequenced on two Illumina NovaSeq 6000 instruments with paired-end sequencing (26×98 bp) at GenomeScan (Leiden, the Netherlands).

### Single-cell RNA sequencing data analysis

Fastq files were mapped to the human (GRCh38) or the mouse (mm10) genome using CellRanger 7.1 (10x Genomics). Filtered matrices were analyzed using Seurat 5.2^39^ in R studio with R 4.4.2. Quality control steps included removing features detected in less than 3 cells and removing cells with less than 200 features, more than 6000 genes and more than 20% mitochondrial RNA. Data was normalized by scaling feature expression by the total expression per cell, multiplied by a scale factor of 10,000 and then log-transformed. This was followed by selection of features that exhibit high cell-to-cell variation and data was linearly transformed prior to dimensionality reduction. Data was subjected to principal component analysis and unsupervised clustering by the Louvain clustering method. The number of PCAs to apply were determined using an ElbowPlot in Seurat. Samples were integrated using Canonical Correlation Analysis (CCA). Cell clusters were visualized using Uniform Manifold Approximation and Projection (UMAP) and were manually annotated using published markers of immune cell subsets. Cluster markers or within each cluster for *Kdm6a^WT^ vs Kdm6a^Δ-Haem^* comparison were generated using the “FindMarkers” function. Gene ontology analysis was performed using the cluster profiler package (Version 4.14.4). Volcano plots were generated using EnhancedVolcano in R (v.1.24.0). Upset plots were generated with UpSetR (v1.4.0).

### Single-cell ATAC sequencing

The nuclei suspensions were counted manually and diluted according to manufacturer’s protocol to obtain 10.000 single cell data points per sample. Each sample was run separately on a lane in Chromium controller with Chromium Next GEM Single Cell ATAC Reagent Kits v1.1 (10xGenomics). Single cell ATACseq library preparation was done using standard protocol. Sequencing was done on Nextseq2000.

### Single-cell ATAC sequencing data analysis

Raw reads were aligned against the mouse genome (mm39, Ensembl 104 release) and counted by StarSolo^40^. Peak calling was done by MACS^41^ and the parameters used were --nomodel --shift -100 --extsize 200 --broad. Preprocessed counts were further analyzed using Scanpy^42^. Basic cell quality control was conducted by the following criteria: a.) mean fragment length = 300, b.) 7.8 < total counts per cell (log) < 12, c.) Reads in peaks > 0.2, d.) Reads in Chr.M < 0.1. Downstream analyzed using peak vs cells matrices were analyzed using Signac^43^ (1.14.0) in R studio with R 4.4.2. Briefly, data were normalized *via* frequency-inverse document frequency (TF-IDF) normalization. Top variable features were selected for dimensional reduction with singular value decomposition (SVD) on the TD-IDF matrix for latent semantic indexing (LSI). The first LSI component was removed from downstream analysis as it was correlated with sequencing depth (visualized with the DepthCor() function in Signac). Cell clusters were then visualized in a low-dimensional space with UMAP. A gene activity matrix was computed by summing the fragments intersecting the gene body and promoter region to visualize canonical markers and interpret cell clusters, and subsequently log-normalized. To additionally help with the deconvolution of cell clusters, we used our unpaired single-cell RNA sequencing data to identify correlation patterns between the gene activity ma trix and scRNA-seq data. This was achieved by identifying a set of anchors between the RNA and ATAC data followed by a transfer of cell labels to the ATAC data. Gene activity was then visualized with CoveragePlot() to ensure validity of cell type assignment. Cluster specific peaks or within each cluster for *Kdm6a^WT^ vs Kdm6a^Δ-Haem^* comparison were generated using the “FindMarkers” function. ATAC-peaks were linked to genes for each cell cluster with the gABC-scoring function from STARE (v.1.0.4)^44^. The unified ATAC peaks were used as candidate enhancer and the counts per million (CPM) in each peak of each cell cluster used as enhancer activity. Scanpy was used for file access^42^. As gene annotation, the M27 version from GENCODE was used^45^. The window size for STARE was set to 5MB, the score cutoff to 0.02 and as contact the estimate based on distance was chosen as follows: “STARE_ABCpp -a gencode.vM27.annotation.gtf.gz -b <unified ATAC peaks> -n 5+ -w 5000000 -f False -o <output_dir>”. Processing involving genomic coordinates made use of pybedtools (0.8.1)^46^. For analysis of the signal coverage, the bam-files for each cell cluster were created with the help of the bam-merge-master functionality from STITCHIT^47^, and subsequently converted to bigwig files with bamCoverage from deeptools (v3.5.1) (normalize with CPM and bin size of 20)^48^. The coverage plots were then generated with the plotHeatmap function from deeptools. For determining the location of regions with respect to genes, each base pair was assigned to a location and the percentage given in relation to the summed base pairs of all peaks. Promoters were defined as ±200 bp around all annotated TSS of all genes, and introns were defined as parts of the gene body that were not labelled as anything else. Colours for the locations were taken from the colorcet Python package based on Glasbey *et al*^49^.Volcano plots were generated using EnhancedVolcano in R (v.1.24.0). Upset plots were generated with UpSetR (v1.4.0).

### Transcription factor footprinting analysis with TOBIAS

For the transcription factor footprinting analysis, the TOBIAS tool was utilised^50^. The scATAC data was pre-processed by the scframework^51^ derived standardised Jupyter notebooks, and cells were grouped according to their cell type. These were then separated by condition (WT/KO) to cell barcodes, and the data was summarised as pseudobulks from the global BAM file, in order to cope with the sparseness of single cells. Per cell type, the two pseudobulk bam files representing the conditions were fed to TOBIAS by making use of the TOBIAS snakemake pipeline. For processing, the following were used: ensemble mm10 release 104 blacklisted regions, fasta sequences and respective gtf file, as well as the Jaspar core vertebrate database from 2024, 10th release^52^.

### Epigenetic Landscape In Silico deletion Analysis (LISA)

Differentially expressed genes identified in the murine cardiac CD45+ single cell RNA-seq data were uploaded to LISA (http://lisa.cistrome.org/) to identify upstream chromatin regulators of differentially expressed genes. Output consisted of adjusted p values. Inferred chromatin regulators data for CCR2+ macrophages and neutrophils were visualized according to significance in Supplementary Figure 3.

### sc-RNAseq/sn-RNAseq integration & cell communication analyses

Integration of scRNA-seq and snRNA-seq data for cell-cell communication analyses was performed as previously described.^12^ Briefly, we filtered monocytes via the subset() function of whole PBMC dataset from HF patients with no CH and HF patients with KDM6A CH. To integrate snRNA-seq data of cardiac tissue and scRNA-seq datasets of PBMCs, we followed the integration as described in section “Single-cell RNA sequencing data analysis” with some alterations. To improve integration of data, we utilized Harmony^53^ with removed variables the sample and sequencing technology used. For crosstalk analysis we used CellChat (2.1.2 version) ^10^. We followed the standard tutorial ‘Comparison analysis of multiple datasets using CellChat’ from the CellChat GitHub repository (https://github.com/jinworks/CellChat).

### Peripheral Blood Mononuclear Cells (PBMCs) isolation and monocyte to macrophage differentiation

PBMCs were isolated from buffy coats of healthy donors, obtained from Blutspendedienst Frankfurt, by density gradient centrifugation. Briefly, PBMC-enriched blood was diluted 1:3 with pre-warmed PBS and layered on top of human Pancoll (1.077 g ml−1, PAN Biotech), followed by centrifugation at 800g for 20 minutes at room temperature without brakes applied. The white ring of PBMCs was transferred to a new 50 mL Falcon tube, and PBS was added to bring the total volume to 50 mL. The sample was then centrifuged at 300g for 10 minutes at room temperature, with acceleration and brakes enabled. The supernatant was discarded, and the pellet was resuspended in 50 mL PBS. Platelets were removed by two additional washing steps and centrifugation at 200g for 10 min and the supernatant was discarded. CD14+ monocytes were isolated using magnetic-activated cell sorting (MACS, Miltenyi Biotec) with CD14 MicroBeads UltraPure (Miltenyi) before proceeding to monocyte-to-macrophage differentiation. Monocytes were differentiated to macrophages with RPMI-1640 media supplemented with 10% FBS, 1% P/S, 10mM HEPES, 1mM sodium pyruvate, 20ng/ml M-CSF for 7 days, with medium change every second day, before proceeding to downstream assays. Donors of buffy coats were three healthy aged males (65-70 years old), with no loss of Y chromosome in all three patients detected by digital PCR. One of the donors carried DNMT3A splice site mutation (VAF 5.70%), another donor carried TET2 mutation (VAF 2.22%), third donor did not show mutations in DNMT3A and TET2 detected in collaboration with MLL (Münchner Leukämielabor).

### siRNA knockdown of KDM6A in CD14+ monocyte-derived macrophages

KDM6A was silenced in CD14+ monocyte-derived macrophages with 100 nM ON-TARGETplus Human KDM6A siRNA SMARTPool (Horizon Dharmacon) by using siTran2.0 transfection reagent (Origene) and 100 nM ON-TARGETplus siRNA Non-targeting Pool as siRNA control (Horizon Dharmacon). Briefly, the medium was replaced with 1 ml of fresh complete medium 30 minutes before transfection. Subsequently, the transfection mix was prepared by adding components in the order specified and vortexing the mixture: 92.1 µl of 1X transfection buffer, 5.5 µl of 20µM siRNA, and 2.4 µl of siTran 2.0 siRNA transfection reagent. The mixture was then incubated for 20 minutes. Following incubation, the transfection mix was added onto the cells in a drop-wise manner, and the plate was moved back and forth to evenly distribute the mix. The media was refreshed 6 hours post-transfection. Finally, supernatants from transfected cells were collected 48 hours post-transfection and the cells were analyzed by qPCR.

### Human cardiac fibroblast activation assay

Primary human cardiac fibroblasts (HCF-c) (#C-12375, Promocell) were cultured in Fibroblast Growth Medium 3 (Promocell) according to the manufacturer’s protocol. HCF-c were seeded at a density of 1200 cells/well in 18-well µ-Slides (IBIDI) coated with human fibronectin (Sigma Aldrich, 0.1% solution; 1:200 in water). Next day, in indirect co-culture experiments, fibroblasts were incubated with supernatants of transfected CD14+ monocyte-derived macrophages. The supernatants were collected 48 h after transfection and were stored at -80°C. Macrophages supernatant was mixed with the fibroblast cell medium at a ratio 1:2 (co-culture medium, 20 µl supernatant and 40 µl medium). Fibroblasts were incubated with this co-culture medium for 72 hours; fixed with 4% paraformaldehyde (in DPBS, Thermo Fisher Scientific) and then were used for immunofluorescent staining.

### Neonatal rat cardiomyocyte stress assay - beating frequency and hypertrophy measurements

Neonatal rat cardiomyocytes (NRCM) were isolated from Sprague Dawley P1 and P2 rat pups. Briefly, pregnant female Sprague Dawley rats were obtained from Janvier (Le Genest Saint-Isle, France). Rat pups were sacrificed by cervical dislocation and hearts were transferred into Hank’s buffered saline solution (-Ca^2+^/-Mg^2+^; Gibco) containing 0.2% 2,3-Butanedione monoxime (BDM; Sigma-Aldrich). Hearts were cut in small pieces and dissociated at 37°C with the enzyme mix (Neonatal Heart Dissociation Kit, mouse and rat, Miltenyi Biotec GmbH) followed by tissue homogenization with the genteMACS™ Dissociator (Miltenyi Biotec GmbH; program m_neoheart_01_01) in a C-tube (Miltenyi Biotec GmbH). Cells were transferred to a 70µm strainer (Falcon), centrifuged (80×g, 5 min) and were resuspended in plating medium (DMEM high glucose, M199 EBS; both from BioConcept, 10% horse serum; Thermo Fisher Scientific), 5% fetal calf serum (Gibco), 2 % l-glutamine (Thermo Fisher Scientific) and penicillin/streptomycin (Roche) and incubated for 1 h and 40 min in 10 cm cell culture dishes (Greiner Bio-One GmbH) at 37°C and 5% CO2. Within the incubation time, the fibroblasts attach to the uncoated culture dish and the supernatant contains the cardiomyocytes and was collected. Cells were counted with the Neubauer-Chamber (Carl Roth).

NRCM were seeded in plating medium at a density of 5000 cells/well in 18-well µ-Slides (IBIDI) coated with rat tail collagen type I (Corning; 1:200 in water). One day after isolation, the medium of cultured NRCM was changed to maintenance medium (DMEM high glucose, M199 EBS (both without l-glutamine by BioConcept), 1% horse serum, 2% l-glutamine and penicillin–streptomycin). For indirect co-culture experiments, NRCM were incubated with supernatants of transfected CD14+ monocyte-derived macrophages. Macrophage supernatant was mixed with the maintanance medium at a ratio 1:2 (co-culture medium, 20 µl supernatant and 40 µl medium). NRCM were incubated with this co-culture medium for 48 hours (beating frequency measurement) and 72 hours (hypertrophy assay).

The cardiomyocyte beating frequency was determined 48 hours after NRCM plating by using a bright-field microscope videos (Zeiss) and counting the number of beats per minute (bpm). For hypertrophic cell size assessment, cardiomyocytes were fixed with 4% paraformaldehyde (in DPBS, Thermo Fisher Scientific) for immunofluorescent staining.

### Immunofluorescence staining for cell culture

After indirect co-culture, cells were washed two times with DBPS and fixed with 4% PFA (Thermo Fisher Scientific) for 10 minutes at room temperature. After two washing steps, each for 5 minutes, with DPBS, cells were permeabilized with 0.1 % Triton X-100 (Thermo Fisher Scientific) in DPBS for 10 minutes. Cells were blocked with 5% donkey serum (abcam) in DPBS for 60 minutes at RT. Primary antibodies were incubated in the same blocking solution overnight at 4 °C. HCF-c were stained with phalloidin (1:400, Thermo Fisher Scientific) and rabbit anti-collagen type 1A1 (1:200, Cell Signaling Technology). NRCM were stained with phalloidin (1:400, Thermo Fisher Scientific) and mouse anti-α-actinin (sarcomeric) antibody (1:200, Thermo Fisher Scientific). Cells were washed 4 times with DPBS, each 5 minutes, and secondary antibodies in DPBS were incubated in for 1 hour at RT. Following dyes and secondary antibodies were used: for HCF-c - DAPI (1:1000, Merck) and donkey anti-rabbit-647 (1:200, Thermo Fisher Scientific), for NRCM - DAPI (1:1000, Merck) and donkey anti-mouse-647 (1:200, Thermo Fisher Scientific). Cells were mounted with Fluoromount-G, Invitrogen). Images were taken using Leica STELLARIS confocal microscope. Z-Stack images were acquired with a resolution of 1024x1024 pixels, a speed of 600 frames per second, line average 2, and a Z-Stack size of approximately 1 µm. Quantification was done using Volocity software version 6.5 (Quorum Technologies).

### Statistical analysis

Statistical analyses were performed using Prism 10 (GraphPad Software Inc.). When Shapiro-Wilk-Test normality test indicated a Gaussian distribution, an unpaired t-test was used, as indicated. Otherwise, Mann-Whitney tests were used. Data are represented as mean ± S.E.M and a p-value < 0.05 was considered significant. For single-cell sequencing RNA experiments a Wilcoxon rank sum test with subsequent Bonferroni correction was used to identify differential genes. For single-cell ATAC experiments a logistic regression test with subsequent Bonferroni correction was used to identify differential peaks.

---

**Supplemental Figure 1.**
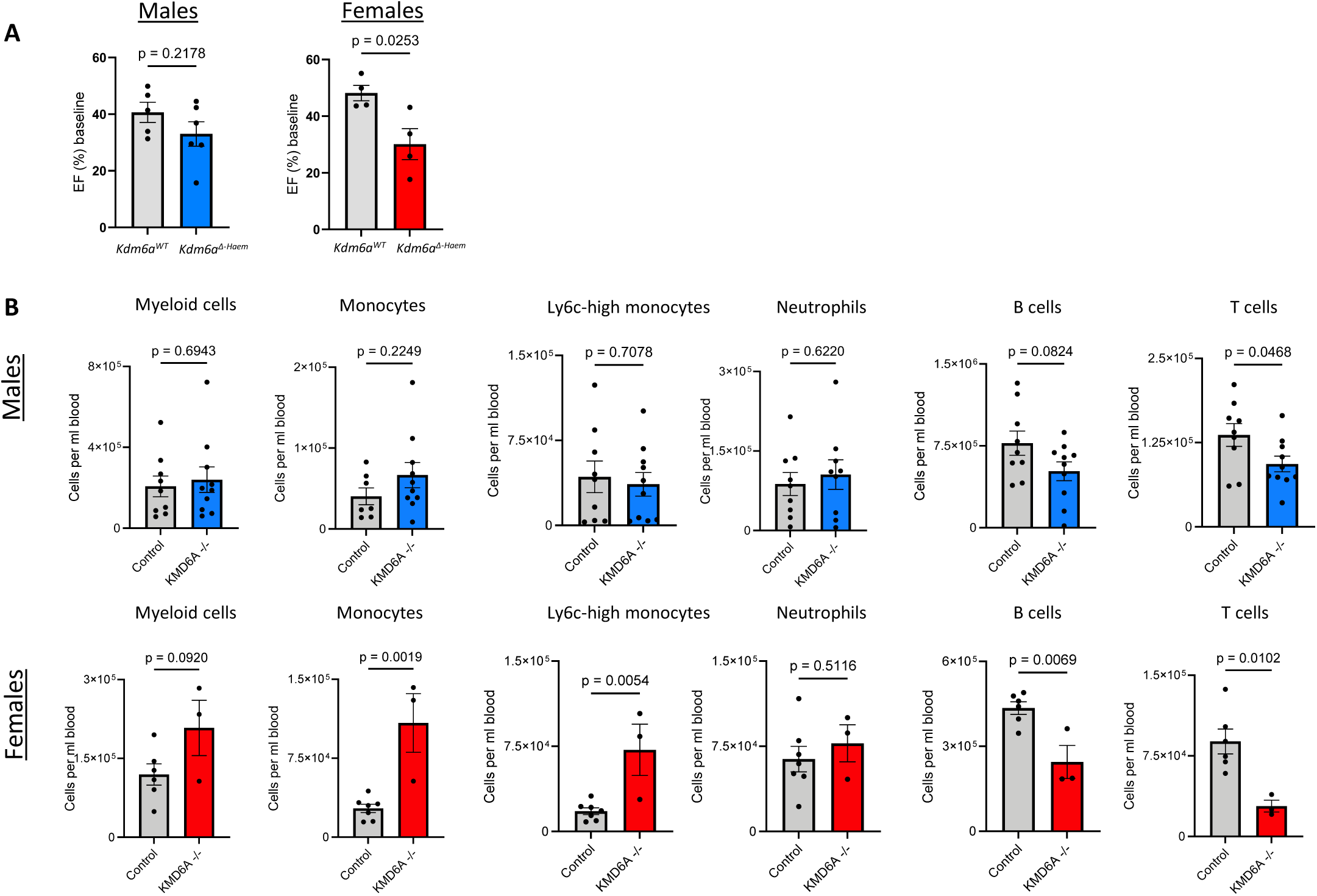
Echocardiography and peripheral leukocyte analysis of Kdm6a^Δ-Haem^ mice, split by sex. (**A**) Cardiac ejection fraction (EF) 28 days post-myocardial infarction (MI) for male (*Kdm6a^WT^* n=5 vs *Kdm6a^Δ-Haem^* n=6, p=0.2178) and female *Kdm6a^Δ-Haem^* mice (*Kdm6a^WT^* n=4 vs *Kdm6a^Δ-Haem^* n=4, p=0.0253) (**B**) Flow cytometric analysis of blood leukocytes of male and female *Kdm6a^Δ-Haem^* mice at 3 days post-MI. Data are represented as mean ± SEM, p-values were calculated using two-tailed unpaired t-tests.

**Supplemental Figure 2.**
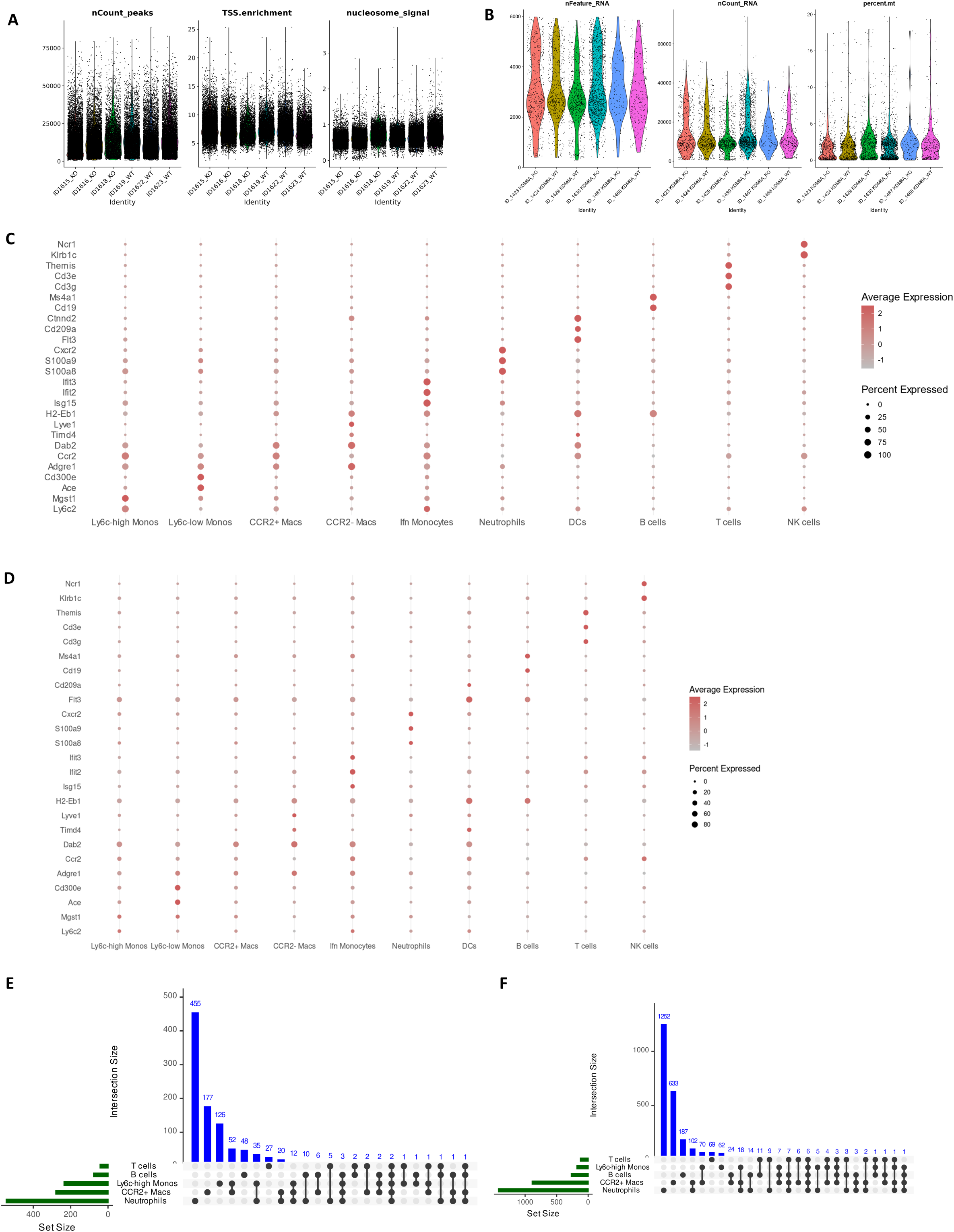
Quality control and annotation of murine scATAC-seq and scRNA-seq data. (**A**) Quality control metrics for filtered scATAC-seq data (**B**) Quality control metrics for filtered scRNA-seq data (**C**) RNA expression of key markers used for annotation of leukocyte subsets in scRNA-seq data (**D**) Gene activity for markers used for scRNA-seq annotation, as assessed by the chromatin accessibility associated with the respective gene (**E**) Intersection plot per cell type of differentially expressed genes from scRNA-seq data (**F**) Intersection plot per cell type of differentially accessible peaks from scATAC-seq data.

**Supplemental Figure 3.**
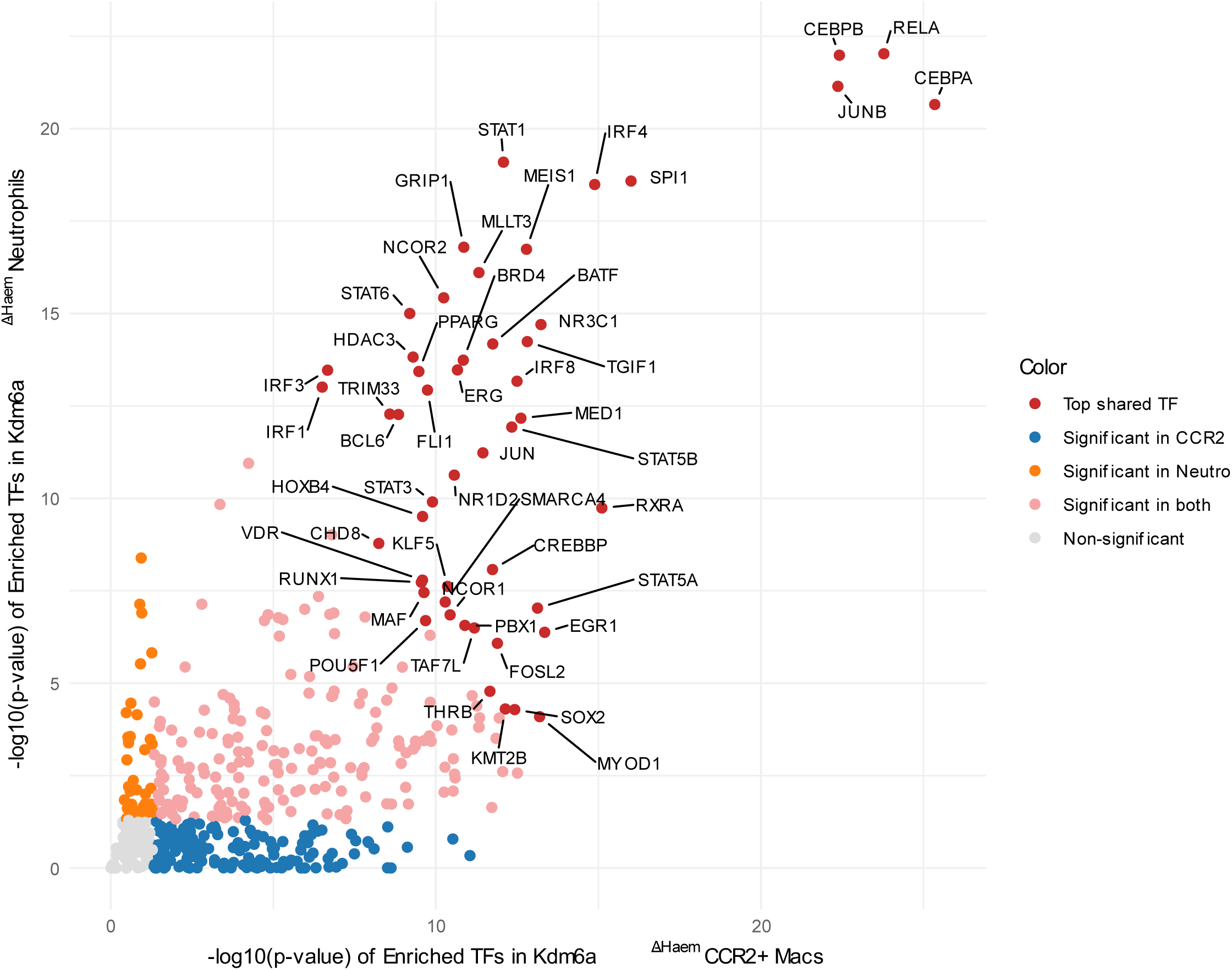
LISA analysis. Differentially upregulated genes from *Kdm6a^Δ-Haem^* neutrophils and CCR2 recruited cells (CCR2 macrophages and Ly6c-high monocytes combined) were used for epigenetic Landscape In Silico deletion Analysis (LISA) to infer upstream transcription factors and/or chromatin regulators responsible for the perturbation of the input genes. Output per cell type was plotted according to significance.

**Supplemental Figure 4.**
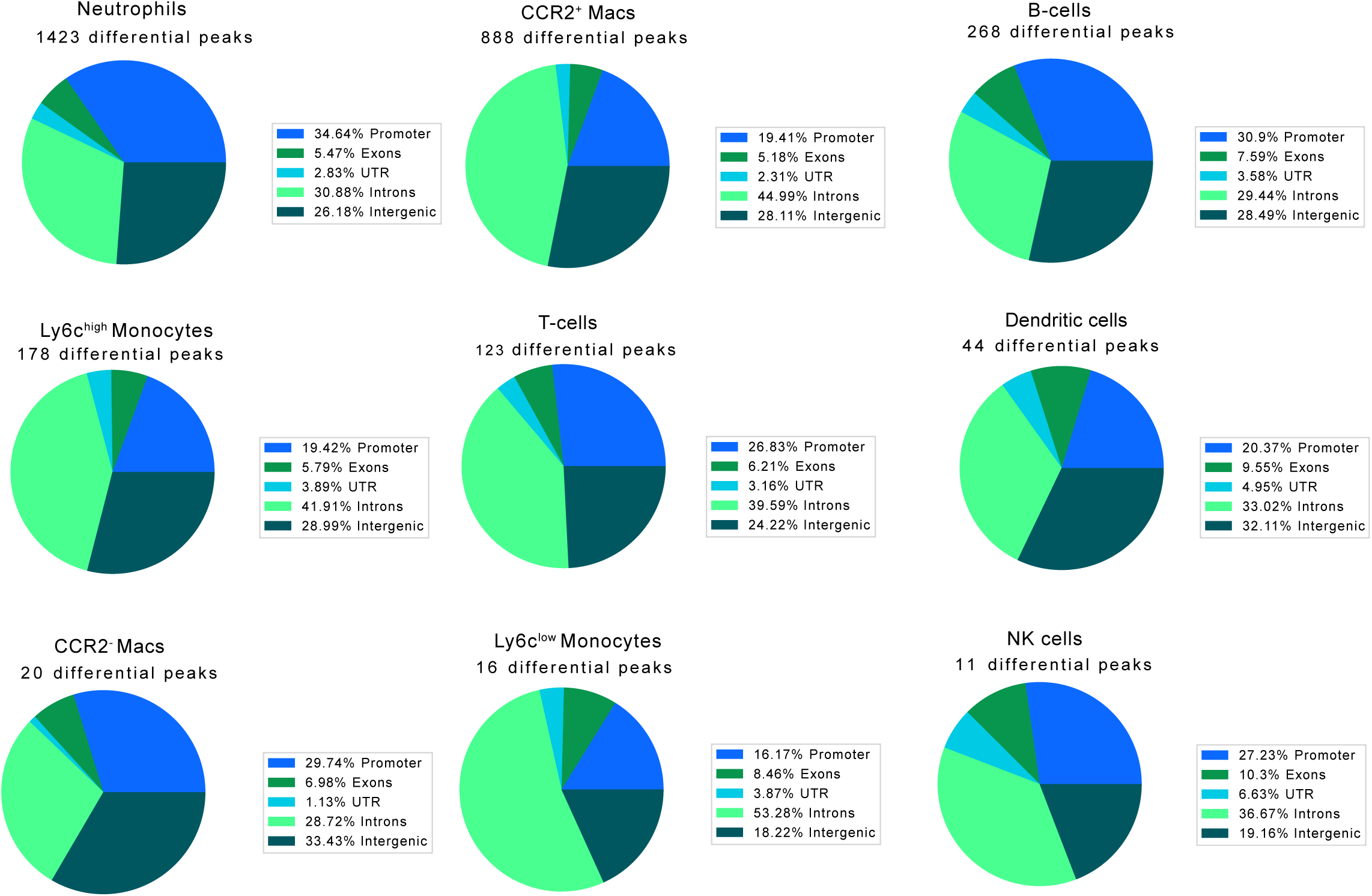
Differential peaks genomic location in Kdm6a^Δ-Haem^ mice. Genomic distribution of differentially accessible chromatin regions across immune cell subsets in *Kdm6a^Δ-Haem^* mice. Each chart indicates the percentage of the differential peaks’ base pairs located within promoters, exons, untranslated regions (UTRs), introns, and intergenic regions. The number of differentially accessible peaks per cell type is indicated above each chart.

**Supplemental Figure 5.**
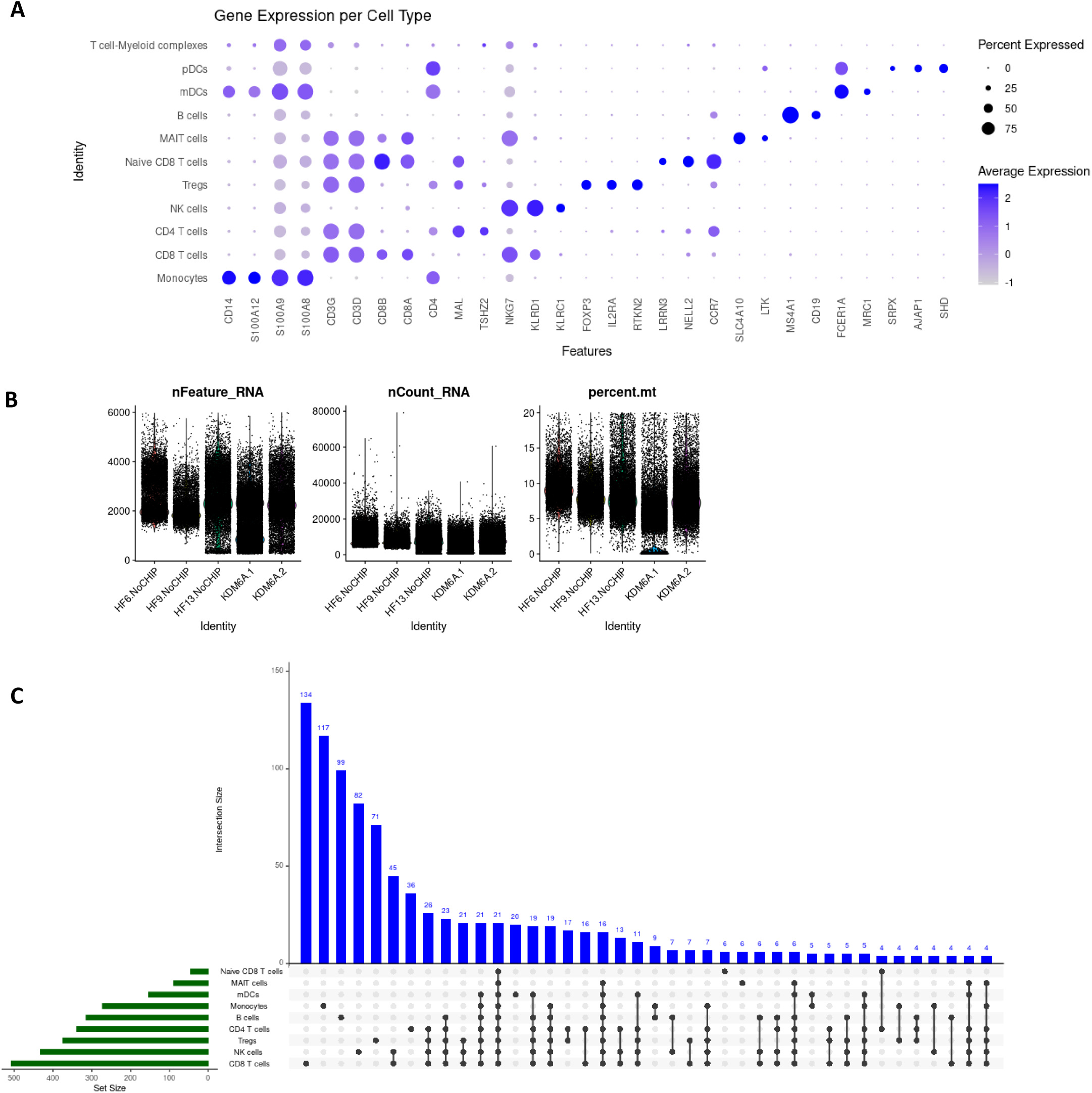
Single cell RNA-seq analysis of PBMC data from HF CH patients. (**A**) RNA expression of key markers used for annotation of peripheral leukocyte subsets. (**B**) Quality control metrics for single-cell RNA-seq data; genes/cells, UMIs/cells, %mitochondrial RNA/cell (**C**) Intersection plot per cell type of differentially expressed genes in HF patients with KDM6A CH.

**Supplemental Figure 6.**
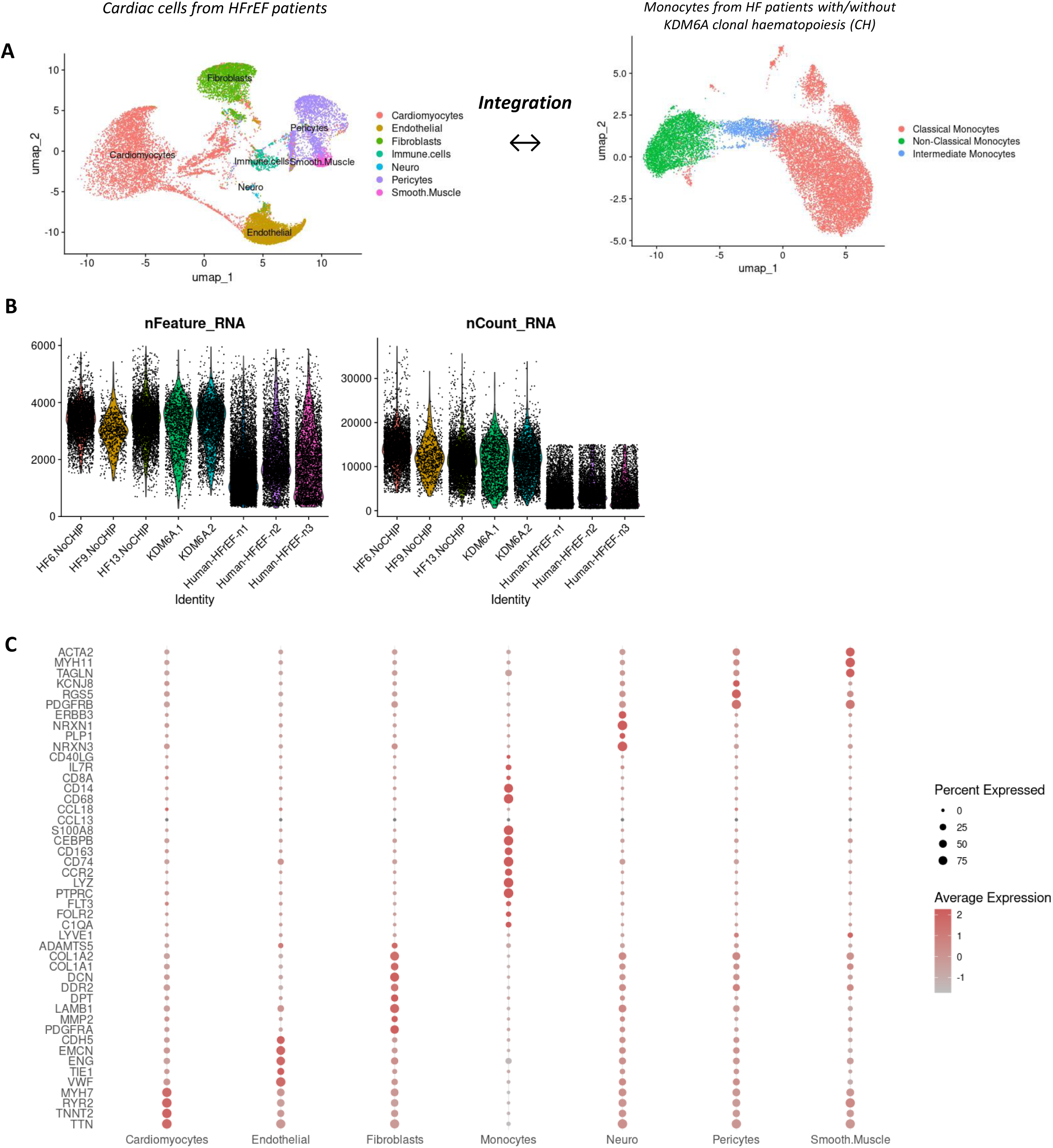
Integration of snRNAseq and scRNAseq data. (**A**) Uniform manifold approximation and projection (UMAP) clustering of single-cell RNA sequencing data from published data from heart failure patients with reduced ejection fraction (HFrEF) and UMAP clustering of subclustered monocytes of HF patients with/without KDM6A clonal haematopoiesis (CH) (**B**) Quality control metrics for filtered single-cell RNA-seq data; genes/cells, UMIs/cells, %mitochondrial RNA/cell (**C**) RNA expression of key markers used for annotation of leukocyte subsets.

